# Brain Kappa Opioid Receptor Availability Across Stress and Social Buffering Conditions: A Positron Emission Tomography Study in Coppery Titi Monkeys

**DOI:** 10.64898/2026.02.17.706461

**Authors:** Claudia Manca, John P. Paulus, Alita J. D Almeida, Anelise Caceres, Meghan J. Sosnowski, Brad A. Hobson, Emilio Ferrer, Abhijit J. Chaudhari, Karen L. Bales

## Abstract

Social connectedness strongly influences health and longevity, and adult pair bonds provide psychological benefits distinct from other social relationships. Oxytocin (OT), corticotropin-releasing hormone (CRH), and opioids, play an important role in the formation and maintenance of pair bonds. Evidence suggests that OT modulates the stress response via the hypothalamic-pituitary-adrenal (HPA) axis, while the kappa (κ) opioid system interacts with and may modulate OT signaling in contexts of stress and separation. In this study 20 coppery titi monkeys were exposed to a physical stressor under three social conditions: baseline (no stressor, partner present), stress (stressor, no partner present) and buffering (stressor, partner present). We predicted stress-induced dynorphin release would reduce κ-opioid receptor availability measured via [¹¹C]GR103545 Positron Emission Tomography (PET) and lower cerebrospinal fluid (CSF) OT, whereas partner presence would mitigate dynorphin release and increase CSF OT, with reduced dynorphin inferred from higher κ-opioid receptor radioligand binding. Our results show condition-dependent differences in [¹¹C]GR103545 binding in several brain regions, including the amygdala and hippocampus, with altered binding in both the stress and social buffering conditions. Cortisol levels were elevated in the stress condition compared to baseline. Females exhibited lower CSF OT levels during stress than at baseline, whereas plasma OT levels did not differ across conditions or between sexes. Spearman correlations revealed no significant associations between plasma and CSF OT. Together, these findings highlight the complex interaction between κ-opioid signaling, OT, and HPA axis activity in the context of social relationships and highlight neuroendocrine mechanisms underlying stress regulation in pair-bonded species.

## Introduction

Social relationships play a fundamental role in the lives of humans and other animals. The protective ability of social bonds to buffer different negative health is an emerging area of research (Donovan et al., 2018). These protective effects have been coined under the social buffering effect and can vary by strength of the bond (i.e. partner versus parent), age, and sex (Hajek et al., 2016; Jarnecke & South, 2014; Kim et al., 2017). Although previous studies have provided multiple examples of how social relationships and interactions may buffer stressors and adverse health outcomes, the underlying neurobiological mechanisms still remain under-explored (Donovan et al., 2018; Kiyokawa & Hennessy, 2018). While there are different types of social relationships, as adults one of our closest relationships is with a romantic partner. Expanding upon Bowlby’s (1969) attachment theory, Hazan and Shaver (1987) suggested that the emotional connections between romantic partners were similar to those found between infants and caregivers, indicating that adult romantic relationships fulfill attachment needs.

Adult pair bonds can provide benefits that are not found in other social relationships (Zeifman, 2019). These attachment relationships share unique psychological features such as distress at separation, social buffering, preference for the familiar partner, and desire for proximity to each other (Hazan & Shaver, 2007; Hennessy et al., 1995; Sbarra & Hazan, 2008). Oxytocin (OT) has gathered particular interest in the context of social buffering as it facilitates stress regulation through its effects on the hypothalamic-pituitary-adrenal (HPA). OT is an evolutionarily conserved neuropeptide hormone involved in mammalian reproduction and implicated in the regulation of mammalian social behavior (Campbell, 2008; Declerck et al., 2010). OT is released both peripherally and centrally in response to diverse psychogenic and physical stressors, playing a significant role as a modulator of the neuroendocrine stress response mediated by the HPA axis (Love, 2018; Neumann et al., 2000; Smith & Wang, 2014).

Social buffering diminishes the cortisol response by reducing the activity of the HPA axis and has been shown to increase peripheral OT levels (Grewen et al., 2005; Pohl et al., 2019). In coppery titi monkeys (*Plecturocebus cupreus*), an OT antagonist treatment blocked fathers’ ability to buffer daughters’ cortisol response, suggesting that blockade of OT receptors can inhibit fathers’ stress-buffering effects (Witczak et al., 2023). Currently, the OT system appears to be a key biological system that underlies the stress buffering mechanism (Hostinar et al., 2014; Sicorello et al., 2020), and there is accumulating evidence that pair mates may moderate the perception of a stressor and the reactivity of the HPA axis (Crockford et al., 2018). Therefore, OT appears to lower levels of corticotropin-releasing hormone (CRH), adrenocorticotropin hormone (ACTH), and cortisol, though the exact pathways by which this happens are not yet fully understood (Crockford et al., 2018; Love, 2018; Wu, 2021).

We know that the κ opioid system and its endogenous ligand, dynorphin, interact closely with CRH and the HPA axis (Bruchas et al., 2010). In animal models of depression, anxiety, and drug-seeking behaviors, it has been demonstrated that activation of the dynorphin/κ system is necessary and sufficient for stress-induced behavioral responses (Bruchas et al., 2010). Dynorphin has been strongly implicated in mediating stress responses, producing feelings of anxiety and dysphoria (Kaski et al., 2021; Knoll & Carlezon, 2010), therefore the κ opioid system is a strong candidate mechanism for the negative affect associated with partner separation. Activation of the κ opioid system in rats suppresses social behavior, whereas blocking κ opioid receptors (KORs) negates the inhibition of social play in an unfamiliar environment (Vanderschuren et al., 1995; Winter & Jurek, 2019).

We have hypothesized that the κ opioid system interacts with and potentially modulates OT in negative aspects of separation and partner loss (Bales & Rogers, 2022). KORs are found on OT neurons in the rat hypothalamus and pituitary (Smith & Wise, 2001). There is also evidence that manipulating KORs influences the release of plasma OT. Specifically, KOR agonists decrease plasma OT levels, while KOR antagonists increase plasma OT levels in rats (Van de Heijning et al., 1991; van Wimersma Greidanus et al., 1996). Oxytocinergic and opioidergic system interactions have been observed across multiple species, with endogenous opioids influencing OT release in various social behaviors (Beery & Kaufer, 2014; Ragen et al., 2015a; Ragen et al., 2015b; Sandi & Haller, 2015); however, this has not been directly investigated in pair bonding and in social buffering.

Like humans, socially monogamous titi monkeys can form strong adult pair bonds and provide an excellent model to better understand the neurobiology of separation and social buffering. Titi monkeys show preference for their pair mate over a stranger (Carp et al., 2016), they display distress upon separation from their pair mate (Cubicciotti & Mason, 1976; Mendoza & Mason, 1986), and their pair mate can buffer their stress response to novelty (Hennessy et al., 1995; Mendoza et al., 2000; Ragen et al., 2013). Separation from the pair mate has physiological and behavioral consequences that include an increase in plasma cortisol concentration, as well as increased locomotion and contact calls for the pair mate (Mendoza et al., 2000; Ragen et al., 2012). Studies have been conducted in titi monkeys on the connection between separation distress and opioids, particularly µ opioids and KOR (Ragen et al., 2013; Ragen et al., 2015b).

This study investigated the role of the κ opioid system in titi monkeys and how the presence of a pair mate may influence an individual against an acute stressor. Our hypothesis is that the release of dynorphin during a physical stressor will lead to decreased availability of KORs, measured via [¹¹C]GR103545 positron emission tomography (PET). In contrast, the presence of the pair mate will mitigate the dynorphin release, leading to an increase of KOR availability and therefore a higher [^11^C]GR103545 binding, and result in an increase in CSF OT. We hypothesized that the presence of a mate would mitigate dynorphin release after an acute stressor in the buffering condition resulting in an increased availability of KORs.

PET neuroimaging also provided an exploratory opportunity to investigate other regions that show dynamic changes in KORs. Finally, we measured how the different social conditions affected HPA activity by measuring plasma cortisol. We predicted that the separation from a mate after an acute stressor would increase cortisol concentrations, while the presence of a mate would reduce cortisol concentrations. We also expected the presence of a mate to increase CSF and possibly plasma OT concentrations.

## MATERIAL AND METHODS

### Subjects

Twenty titi monkeys were used as subjects in this study. Females were not on hormonal birth control and were paired with vasectomized males or had received a tubal ligation (one subject). They were born and housed at the California National Primate Research Center (CNPRC). All animals were fed daily at 0800 and 1300 h on a diet consisting of monkey chow, carrots, apples, bananas, and rice cereal. Water was available *ad libitum.* Rooms were maintained at 21°C and they were set on a 12:12-h light:dark cycle with lights on at 0600 and lights off at 1800. The animals were housed in cages measuring 1.2 m x 1.2 m x 2.1 m or 1.2 m x 1.2 m x 1.8 m. These housing conditions were similar to those described in Mendoza & Mason (1986) and Tardif et al. (2006). At the beginning of the experiment, the mean age of the subjects was 6.4 years (range 2.31-15.90) and the mean duration of pairing was 2.3 years (range 0.25 – 5.57). All procedures were approved by the Institutional Animal Care and Use Committee (IACUC; protocol number #23483) of the University of California, Davis.

### Conditions

We used [^11^C]GR103545 PET to examine the effects of the presence and absence of social buffering on endogenous κ receptor availability and CSF OT. This radiotracer has been previously characterized in titi monkeys (Almeida et al., 2025), other non-human primates (Schoultz et al., 2010; Talbot et al., 2005) and humans (Naganawa et al., 2014). This experimental protocol consisted of three within-subject conditions: baseline, stress, and buffering. In the baseline condition, subjects were not exposed to an experimental stressor prior to anesthesia (see PET scan procedure). Subjects were removed from their cage, administered the radiotracer, and immediately positioned in the PET scanner (Figure 1A).

**Figure 1.**
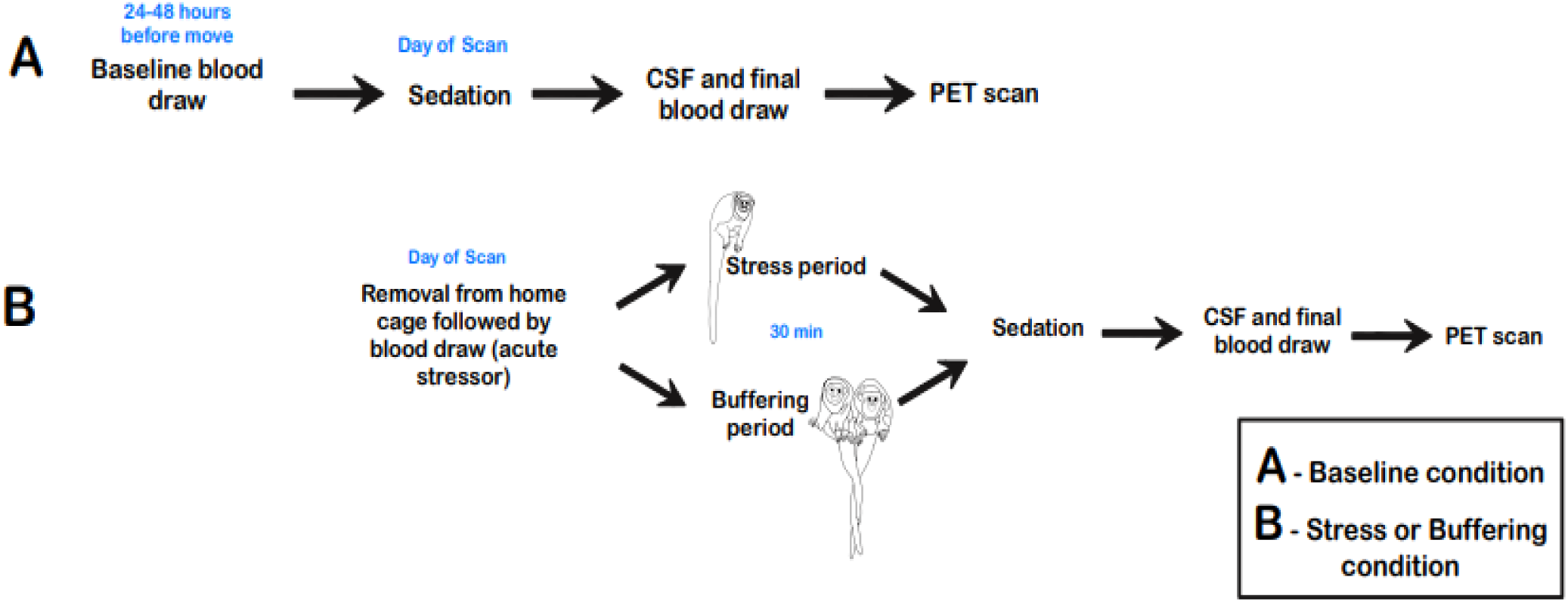
Experimental Timeline

In the stress condition, subjects underwent an acute physical stressor (removal from home cage followed by a first blood draw of 1ml). They were then returned to a transport box (31 × 31 × 33 cm) *without* their pair mate for a minimum of 30 minutes. During that time, the subject’s pair mate was removed from the room and transported to a different room. Following the 30 minute separation period, the subject was removed from the transport box and positioned in the scanner for PET imaging (Figure 1B).

In the buffering condition, the subject underwent the same physical stressor described above (subject was restrained for a 1 ml blood draw). However, following the stressor, the subject was placed into the transport box *with* their pair mate for at least 30 minutes. After this period, the subject was then removed from the transport box and positioned in the scanner for PET imaging (Figure 1B).

Experimental conditions were counterbalanced across subjects with a minimum interval of a month between successive scans (mean interval: 38 days). Subjects spent an average of 42.28 minutes (SD = 5.38) in the transport box prior to PET imaging for both the buffering and stress condition.

### PET Scan Procedures

Positron emission tomography (PET) scans were performed on all subjects three times, once per experimental condition. Forty-eight hours prior to scans, the subject and their pair mate were relocated to a metabolism room to reduce potential confounding of novel housing on brain metabolism, behavior, and endocrine activity. On the day of the PET scan, the subject was removed from the home cage and depending on the social condition, underwent an acute physical stressor (manual restraint for a blood draw) before being prepared for PET scanning. For the stress and social buffering conditions, subjects were transferred to a transport box. They were restrained, and a blood sample was collected before being returned to their transport box either *with (buffering)* or *without (stress)* their pair mate. Following this interval, subjects were manually restrained, and given Ketamine (25 mg/kg IM) for sedation; a second blood draw (1ml) and CSF sample were collected. The blood sample was collected from the femoral vein into a heparinized syringe. Both the blood and the CSF samples were immediately placed on ice. An endotracheal tube was placed, and two intravenous catheters were placed in the saphenous vein of both legs to administer IV fluids (lactated ringers solution, 10 ml/kg/h). The subject was positioned on the bed of a dedicated brain PET scanner (PiPET, Brain Biosciences, Rockville, MD), and the [^11^C]GR103545 radiotracer was administered intravenously. The dynamic 110 min PET acquisition began approximately 15 s prior to injection of [^11^C]GR103545 (injected activity 47.7±5.7 MBq). Data were reconstructed with an isotropic voxel size of 0.8 mm^3^ using framing of 6 × 10 s, 8 × 30 s, 5 × 60 s, 4 × 300 s, 8 × 600 s. A single-pass “cold” transmission scan was acquired pre-injection for attenuation and scatter correction. Anesthesia was maintained throughout the scan with isoflurane (1-2%). Following the scan, animals were housed in a metabolism room until radioactivity had decayed to background levels and then returned to their home cages with their partners.

### MRI scanning

Structural magnetic resonance imaging (MRI) scans were conducted on a different day, in a GE Signa LX 9.1 scanner (General Electric Corporation, Milwaukee, WI, USA) with a 1.5 T field strength and a 3” surface coil. Each subject was fasted 8-12 h before the procedure. At the start of the procedure, the subject was sedated with ketamine (10 mg/kg IM) and midazolam (0.1 mg/kg IM), and an endotracheal tube was placed. The MR acquisition employed a 3D spoiled gradient echo pulse sequence in the coronal plane using the following parameters: echo time (TE) = 7.9 ms, repetition time (TR) = 22.0 ms, flip angle=30.0°, field of view=8 cm, number of excitations = 3, matrix size = 256 × 256, resulting in a pixel size of 0.3125 × 0.3125 mm^2^ and a slice thickness of 1 mm. The subject’s EtCO2, oxygen saturation, blood pressure, and heart rate were monitored throughout the scan. If a subject had undergone an MRI within the same year as the start of this project, that existing MRI was utilized.

### PET and MRI Coregistration and Image Analysis

A standardized image processing pipeline specifically tailored to titi monkey PET and MRI scans was developed. In the pipeline each animal’s anatomical MRI scan was resampled to an isotropic voxel size (0.3mm^3^), mapped onto a titi monkey reference space, skull stripped and bias-corrected. A titi monkey brain atlas was then warped to the MRI scan using the symmetric normalization algorithm with the open-source ANTs toolkit (Avants et al., 2011). Separately, reconstructed PET scans were imported into the PMOD (version 4.4) software where the data were visually examined for quality, motion-corrected, and then manually aligned with the MRI. The atlas labels were then transferred to the PET to obtain region-wise time-activity curves. Region-specific, non-displaceable binding potential (BP_ND_) was calculated using the simplified reference tissue model (SRTM) with the cerebellum as the reference region (Almeida et al., 2025).

### Regions of Interest (ROIs)

Regions of interest (ROIs) in this study included: NAcc, septum, amygdala, hippocampus, hypothalamus. Exploratory regions included: putamen, cingulate cortex, orbitofrontal cortex, ventral pallidum, claustrum, parahippocampal gyrus, thalamus, and insula. The rationale for selecting these regions is provided next.

KORs in the NAcc shell may contribute to pair-bond maintenance in titi monkeys. Comparative research with prairie voles has shown that administering KOR antagonists in this region to a paired male attenuates aggression towards a stranger (Resendez et al., 2012). Work in titi monkeys suggests the lateral septum (LS) may be involved in pair bonding as demonstrated by differences in glucose uptake between paired and unpaired males (Bales et al., 2007, 2017). The LS has long been implicated in the control of stress responses and anxiety (Anthony et al., 2014; Sheehan et al., 2004). The LS also has strong reciprocal connections to the hippocampus, a region involved in memory consolidation (Bales et al., 2017; Sheehan et al., 2004).

The hippocampus and the amygdala are not only involved in emotional responses like fear, anxiety, and arousal, but they also play a role in cognitive processes, such as memory and attention, which can carry an emotional aspect (Escriche Chova et al., 2023; Gallagher & Chiba, 1996; Phelps, 2004). A study in mice showed that upregulation of dynorphin/KOR system in the dorsal hippocampus plays a critical role in formation of aversive emotion associated with morphine withdrawal (Chen et al., 2023). A different study in mice found that stress-induced dysphoria engages the dynorphin/KOR system in the hippocampus, BLA, and NAcc (Land et al., 2008). In rats, KORs in the basolateral amygdala (BLA) have been shown to mediate anxiety (Knoll et al., 2011). Thus, KORs in the BLA in titi monkeys may be activated when anxious as a result of separation from a pair mate.

The hypothalamus processes signals from various brain areas to modulate many endocrine systems, including OT, and it is a region of interest particularly for the role of the PVN in activating the HPA axis. Research in rats has shown that KORs are expressed on OT magnocellular neurons, as well as on their projections and terminals in the PVN, providing a pathway through which opioid agonists can directly inhibit OT release (Smith & Wise, 2001b).

Prior work in titi monkeys using receptor autoradiography showed that all of these regions express KORs, with no detectable sex differences in their distribution, although this conclusion was based on a small sample (3 females, 6 males) (Ragen et al., 2015b).

Exploratory regions were chosen because they broadly participate in networks that integrate stress buffering, emotion, social reward, motivation, and the evaluation of social and environmental cues (Blumenthal & Young, 2023; Burkett et al., 2011; Chun et al., 2022; Kim et al., 2016; Lamm et al., 2011; Liu et al., 2019; Niu et al., 2022; Peen et al., 2021).

### Blood Sampling and Hormone Analysis

Blood and CSF samples were collected immediately after anesthesia for the PET scan and placed on ice. Blood samples were collected at a mean of 3.39 min (SD = 1.82) after capture of the subject for the PET scan, and CSF samples were collected at a mean of 9.85 min (SD = 2.42) after capture. Blood samples were centrifuged at 3000 RPM for 15 min at 4°C. Plasma was separated and transferred into tubes, and plasma and CSF samples were stored at -80°C until assay. CSF samples were assayed for OT. Plasma samples were assayed for cortisol and OT.

We used a commercially available enzyme immunoassay kit (ELISA; Arbor Assay, Ann Arbor, MI, USA) CSF samples were not extracted prior to assay, as per previous literature suggesting that CSF samples do not require extraction (Monte et al., 2014; Parker et al., 2010; Tabak et al., 2023). Due to low sample volume, most samples were run without a duplicate, on a single plate. Whenever sample volume permitted, samples were run in duplicate. For these samples (n = 5), the intra-assay co-efficient of variation (CV) was 4.37%. Out of the 60 attempts at collecting CSF samples, 29 samples did not meet the minimum required volume for analysis. 31 remaining samples were included in the final analysis (n_baseline_ = 11, n_buffering_ = 11, n_stress_ = 9).

Plasma samples were extracted prior to assay using a reduction/alkylation procedure, consistent with previous literature indicating that extraction is necessary for plasma to yield valid results (Szeto et al., 2011; Tabak et al., 2023). We used a commercially available enzyme immunoassay kit (ELISA; Arbor Assay, Ann Arbor, MI, USA) that was validated for use with titi monkey CSF and plasma samples (see Supp. Materials A). Samples were run in duplicates. The intra-assay coefficients of variation (CVs) were 5.50%, 5.18%, and 5.13% for plates 1-3, respectively.

Plasma cortisol was assayed at the UC Davis Endocrinology Laboratory. Plasma cortisol concentrations were estimated in duplicate using commercial radioimmunoassay kits (Siemens Healthcare, Malvern, PA, USA), previously validated both chemically and biologically for titi monkeys (see Witczak et al., 2021). A total of five plates were assayed, with intra-assay CVs of 9.9%, 10.3%, 7.3%, 4.8%, and 9.1%, with an inter-assay CV of 4.0%.

### Statistical Analyses

All analyses were conducted in R (R Core Team, 2022) using the *lme4* and *lmerTest* packages. Linear mixed-effects models (LMMs) were fit for cortisol, CSF OT, plasma OT, and PET BP_ND_, with condition and sex as fixed effects and animal ID as a random intercept to account for repeated measures. Models followed the structure *outcome ∼ condition × sex + (1|ID)*, and assumptions of normality and homoscedasticity were evaluated via inspection of residual and Q–Q plots. A singular fit occurred for plasma OT and for the septum, indicating minimal random-effects variance; however, the LMM structure was retained for consistency across outcomes and due to the repeated-measures design. For PET data, separate LMMs were estimated for each region, followed by an ANOVA to test omnibus effects.

After the planned primary analysis (LMM and ANOVA), we conducted exploratory secondary analyses to investigate potential subgroup differences. Simple effects were examined using estimated marginal means (emmeans) to assess (i) differences between experimental conditions and (ii) sex differences within conditions. These contrasts were specified a priori based on the experimental design and previous literature showing sex differences in cortisol and OT levels, and sex differences in the κ opioid system (Kirschbaum et al., 1992; Kramer et al., 2004; Marazziti et al., 2019; Negus et al., 2002; Roberts et al., 1998; Vijay et al., 2016). Thus, we planned to carry out simple effects to test *a priori* hypotheses, even in the absence of statistically significant interactions. FDR correction was applied to exploratory imaging regions but not to regions with *a priori* predictions. Statistical significance was set at α = 0.05. Effect sizes are reported as partial η² for ANOVA terms, marginal and conditional R² for overall model fit, and Hedges’ g (small sample corrected) for pairwise contrasts. All comparisons are reported in the Supplementary Materials (see Supp. Materials B).

## 3. Results

### 3.1 Cortisol

A LMM was conducted to examine the effects of condition and sex on cortisol levels, with subject ID included as a random intercept to account for repeated measures. Random effects indicated substantial between-subject variability (σ² (ID) = 51,695, SD = 227.4), while residual variance was σ² = 20,547 (SD = 143.3).

A Type II ANOVA using Satterthwaite’s method revealed a significant main effect of condition (F(2, 36) = 10.30, **p < .001\*\*\***, ηp² = .36), whereas neither the main effect of sex (F(1, 18) = 0.15, p = .702, ηp² = .01) nor the condition × sex interaction (F(2, 36) = 0.74, p = .483, ηp² = .04) was significant.

Planned simple effects, Tukey-adjusted for multiple comparisons, indicated that, among females, cortisol levels were significantly higher in the stress condition relative to baseline (β = –229.02, SE = 66.58, t(35.29) = –3.44, **p = .0042\***, *g* = –0.58). Among males, cortisol levels were also significantly higher in the stress condition compared to baseline (β = –175.68, SE = 64.11, t(35.05) = –2.74, **p = .0253\***, *g* = –0.32), with no other pairwise differences reaching significance.

Sex differences within each condition were non-significant.

**Table 1:**
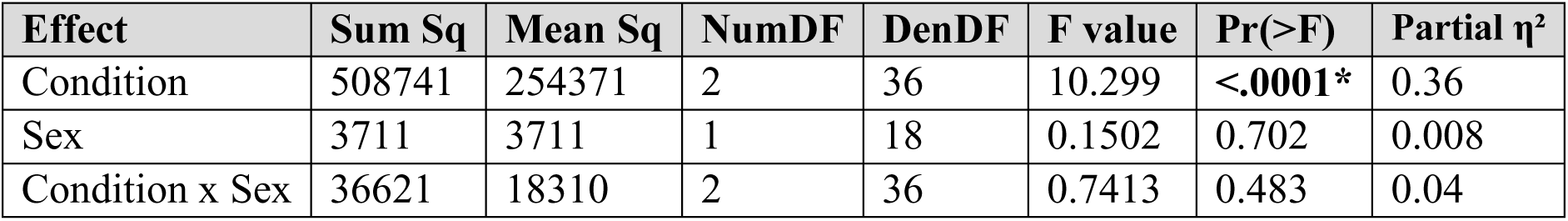
Type II ANOVA results for cortisol concentrations.

**Figure 2:**
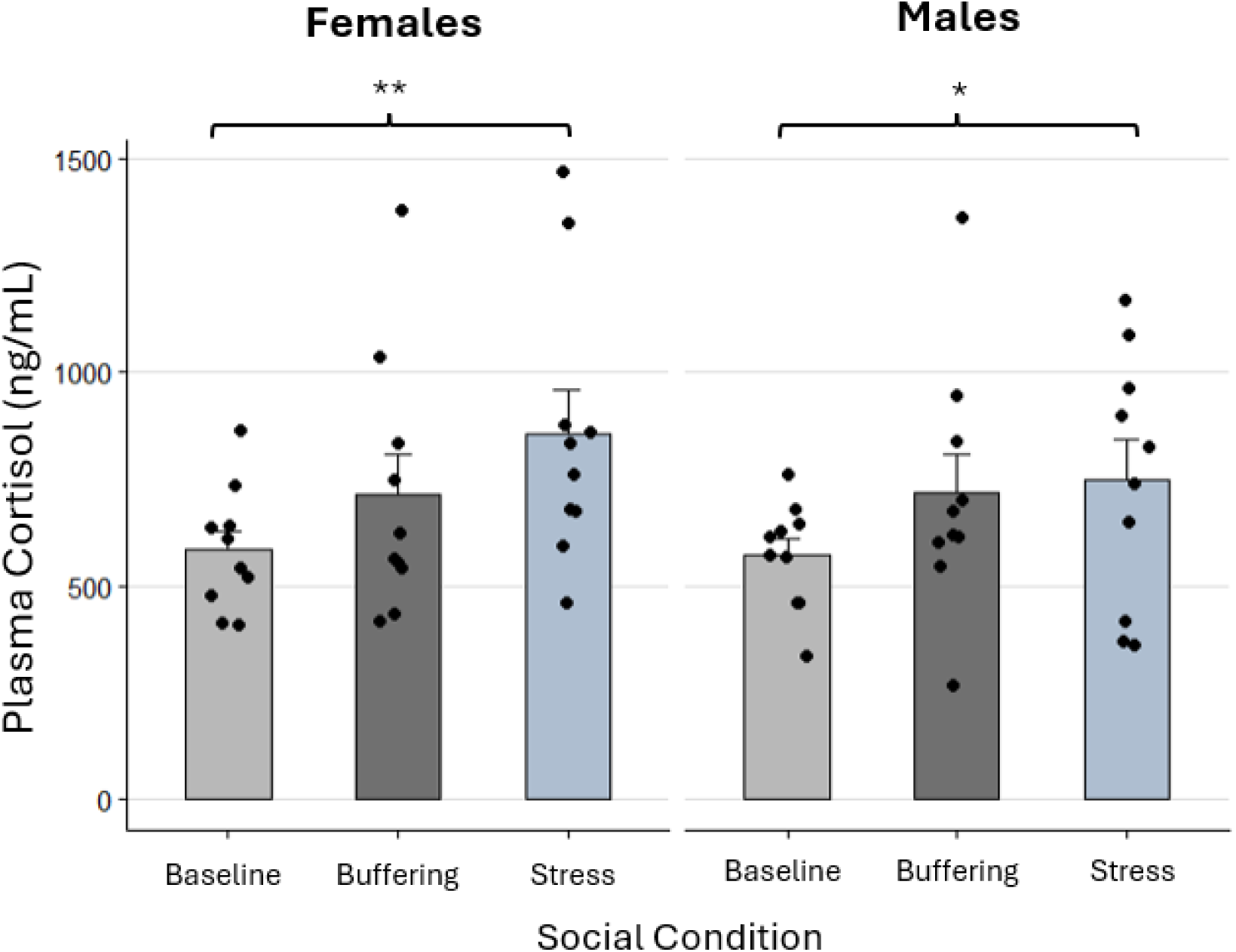
Mean Cortisol Levels by Sex and Condition. Mean cortisol plasma levels are shown for females and males under baseline, buffering, and stress conditions. The error bars represent standard error. Individual data points represent observed values for each subject. Significant differences in cortisol levels were found between conditions (F(2, 36) = 10.30, **p < .001\*\*\***). Simple effects indicated higher cortisol levels in the stress condition compared to baseline in both females (**p = 0.0042\*\***) and males (**p = 0.0253\***), with no significant difference between baseline and buffering. * p<0.05, ** p<0.01.

### 3.2 PET Imaging

#### Amygdala

LMMs were used to examine the effects of condition, sex, and their interaction on amygdala BP_ND_. LMMs results are reported in table 2.

**Table 2.** Results of the linear-mixed effects models of *a priori* regions.

| Brain Region |  | Estimate | s.e | df | t-value | p-value | Marginal R<sup>2</sup> | Conditional R<sup>2</sup> |
| --- | --- | --- | --- | --- | --- | --- | --- | --- |
| Amygdala | Intercept | 1.34348 | 0.0443 | 15.63 | 30.313 | **< 0.001** | 0.124 | 0.556 |
|  | Condition (Buffering) | -0.06374 | 0.0337 | 25.24 | -1.889 | 0.07 |  |  |
|  | Condition (Stress) | 0.01992 | 0.0363 | 26.35 | 0.55 | 0.587 |  |  |
|  | Sex (Male) | -0.03523 | 0.0443 | 15.63 | -0.795 | 0.439 |  |  |
|  | Buffering × Sex (Male) | -0.00351 | 0.0337 | 25.24 | -0.104 | 0.918 |  |  |
|  | Stress × Sex (Male) | -0.07802 | 0.0363 | 26.35 | -2.152 | **0.041\*** |  |  |
| Hippocampus | Intercept | 0.8553 | 0.0243 | 19.55 | 35.14 | **< 0.001** | 0.191 | 0.401 |
|  | Condition (Buffering) | -0.01749 | 0.0247 | 30.27 | -0.707 | 0.485 |  |  |
|  | Condition (Stress) | 0.02231 | 0.0264 | 31.89 | 0.846 | 0.404 |  |  |
|  | Sex (Male) | -0.05847 | 0.0243 | 19.55 | -2.402 | **0.026\*** |  |  |
|  | Buffering × Sex (Male) | 0.01767 | 0.0247 | 30.27 | 0.714 | 0.48 |  |  |
|  | Stress × Sex (Male) | -0.04562 | 0.0264 | 31.89 | -1.729 | 0.093 |  |  |
| Hypothalamus | Intercept | 1.09187 | 0.0781 | 13.36 | 13.975 | **< 0.001** | 0.066 | 0.449 |
|  | Condition (Buffering) | 0.02856 | 0.0662 | 23.11 | 0.431 | 0.67 |  |  |
|  | Condition (Stress) | 0.06033 | 0.071 | 24.51 | 0.85 | 0.404 |  |  |
|  | Sex (Male) | -0.00758 | 0.0781 | 13.36 | -0.097 | 0.924 |  |  |
|  | Buffering × Sex (Male) | 0.12415 | 0.0662 | 23.11 | 1.875 | 0.074 |  |  |
|  | Stress × Sex (Male) | -0.02072 | 0.071 | 24.51 | -0.292 | 0.773 |  |  |
| NAcc | Intercept | 1.58074 | 0.0395 | 16.12 | 40.042 | **< 0.001** | 0.115 | 0.298 |
|  | Condition (Buffering) | -0.05128 | 0.0426 | 27.4 | -1.204 | 0.239 |  |  |
|  | Condition (Stress) | 0.0627 | 0.0454 | 29.36 | 1.383 | 0.177 |  |  |
|  | Sex (Male) | -0.05883 | 0.0395 | 16.12 | -1.490 | 0.155 |  |  |
|  | Buffering × Sex (Male) | -0.01363 | 0.0426 | 27.4 | -0.320 | 0.752 |  |  |
|  | Stress × Sex (Male) | -0.02552 | 0.0454 | 29.36 | -0.563 | 0.578 |  |  |
| Septum | Intercept | 1.44173 | 0.0404 | 44.00 | 35.707 | **< 0.001** | 0.088 | NA singular fit |
|  | Condition (Buffering) | -0.06111 | 0.0557 | 44.00 | -1.096 | 0.279 |  |  |
|  | Condition (Stress) | 0.03549 | 0.0588 | 44.00 | 0.604 | 0.549 |  |  |
|  | Sex (Male) | -0.04927 | 0.0404 | 44.00 | -1.220 | 0.229 |  |  |
|  | Buffering × Sex (Male) | -0.05635 | 0.0557 | 44.00 | -1.011 | 0.318 |  |  |
|  | Stress × Sex (Male) | -0.00128 | 0.0588 | 44.00 | -0.022 | 0.983 |  |  |
| The model (BP<sub>ND</sub> ~ condition × sex + (1\|ID) column indicated the statistical model tested. Bolded values indicate the *p* value was significant at *p* < 0.05. All models included subject as a random effect. For this table, s.e. indicated the standard error of the corresponding parameter estimate. df indicates the degrees of freedom. |  |  |  |  |  |  |  |  |

**Figure 3:**
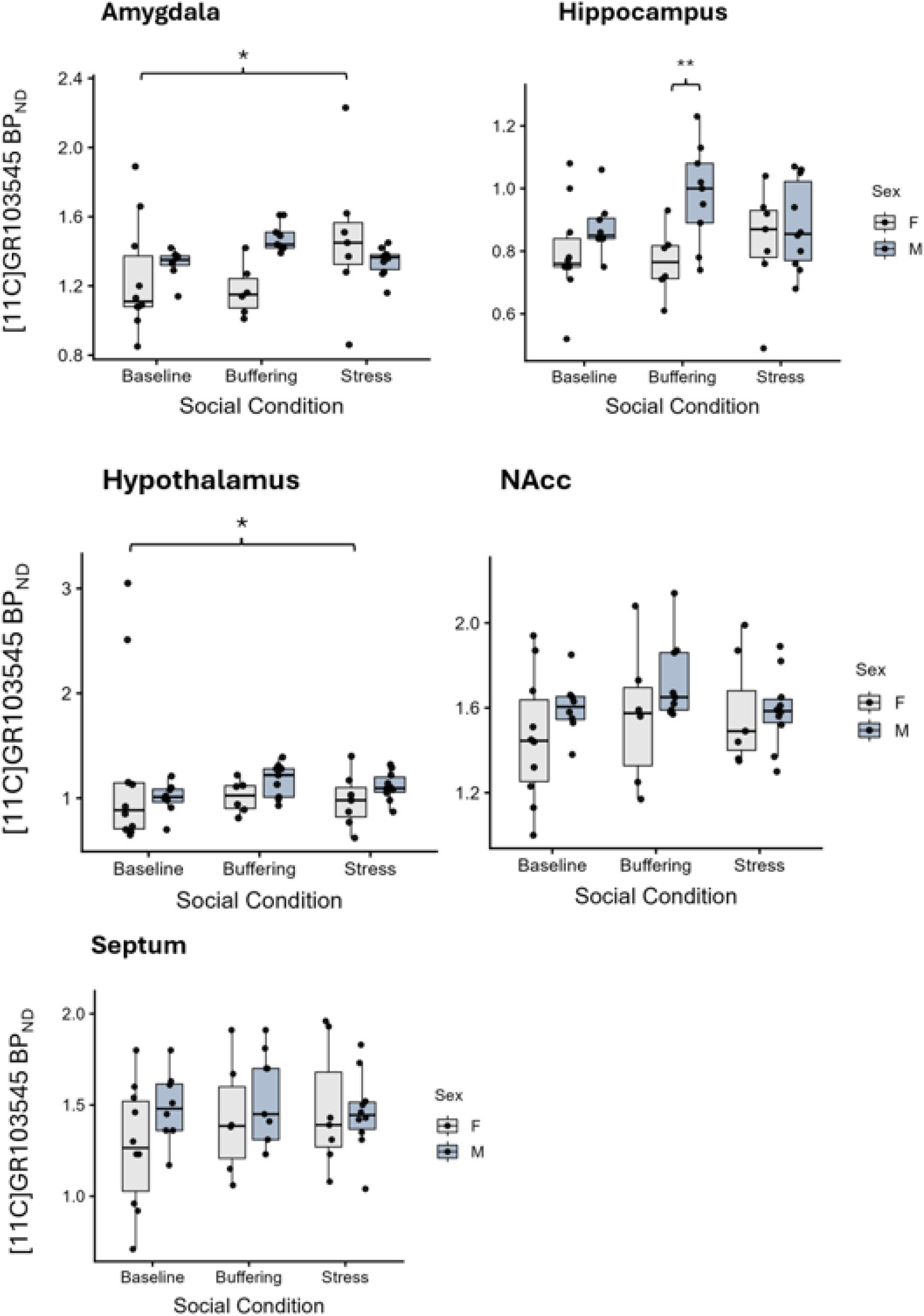
Regional KOR BP_ND_ across Sex and Social Conditions. Boxplot show [11C]GR103545 values for females (light gray) and males (light blue) in the baseline, buffering, and stress conditions across five ROIs. Boxes indicate the interquartile range with the median shown; whiskers extend to 1.5 x the interquartile range. Individual data points are overlaid. In the amygdala, females exhibited significantly higher BP_ND_ in the stress condition compared to baseline (**p = 0.032\***). In the hippocampus, males showed significantly higher BP_ND_ in the buffering condition compared to females (**p = 0.008\*\***). In the hypothalamus, females showed significantly higher BP_ND_ in the baseline condition compared to stress (**p = 0.048\***). No significant condition or sex differences were observed in the NAcc or septum.

Type III ANOVA revealed no significant main effect of condition (F(2, 25.98 = 1.89, p = 0.171, partial η² = 0.13) or sex (F(1, 15.63) = 0.63, p = 0.439, partial η² = 0.039), but marginally significant interaction between condition and sex (F(2, 25.98) = 3.25, p = 0.055, partial η² = 0.20).

Exploratory comparisons showed that in females, BP_ND_ in the stress condition was significantly higher than baseline (estimate = –0.1926, SE = 0.0852, df = 28.3, t = –2.261, **p = 0.032*,** Hedges’ g = –0.43). The buffering–stress comparison was marginal (estimate = –0.1835, SE = 0.1001, df = 30.1, t = –1.833, p = 0.077, g = –0.33), and the baseline–buffering comparison was not significant in females. In males, no within-sex contrasts were significant.

#### Hippocampus

LMMs were used to examine the effects of condition, sex, and their interaction on hippocampus BP_ND_. LMMs results are reported in table 2.

Type III ANOVA revealed no significant main effect of condition (F(2, 31.30) = 0.41, p = 0.667, partial η² = 0.026). The main effect of sex was significant (F(1, 19.55) = 5.77, **p = 0.026\***, partial η² = 0.228). The interaction between condition and sex was not significant (F(2, 31.30) = 1.51, p = 0.236, partial η² = 0.088).

Exploratory comparisons indicated a significant sex difference in BP_ND_ in the buffering condition (estimate = –0.2082, SE = 0.0750, df = 42.8, t = –2.776, **p = 0.008\*\***, g = –0.85). No other comparisons between experimental conditions revealed any significant differences.

#### Hypothalamus

LMMs were used to examine the effects of condition, sex, and their interaction on hypothalamus BP_ND_. LMMs results are reported in table 3.

**Table 3:**
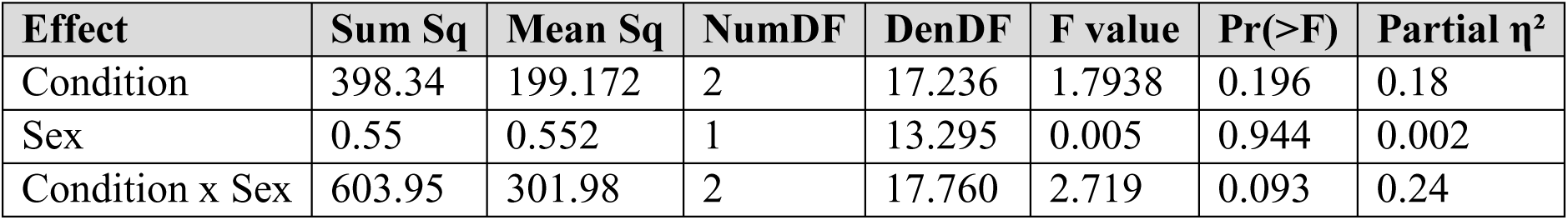
Type II ANOVA results for CSF OT concentrations.

| Effect | Sum Sq | Mean Sq | NumDF | DenDF | F value | Pr(>F) | Partial $\eta^2$ |
| --- | --- | --- | --- | --- | --- | --- | --- |
| Condition | 398.34 | 199.172 | 2 | 17.236 | 1.7938 | 0.196 | 0.18 |
| Sex | 0.55 | 0.552 | 1 | 13.295 | 0.005 | 0.944 | 0.002 |
| Condition x Sex | 603.95 | 301.98 | 2 | 17.760 | 2.719 | 0.093 | 0.24 |

**Figure 4.**
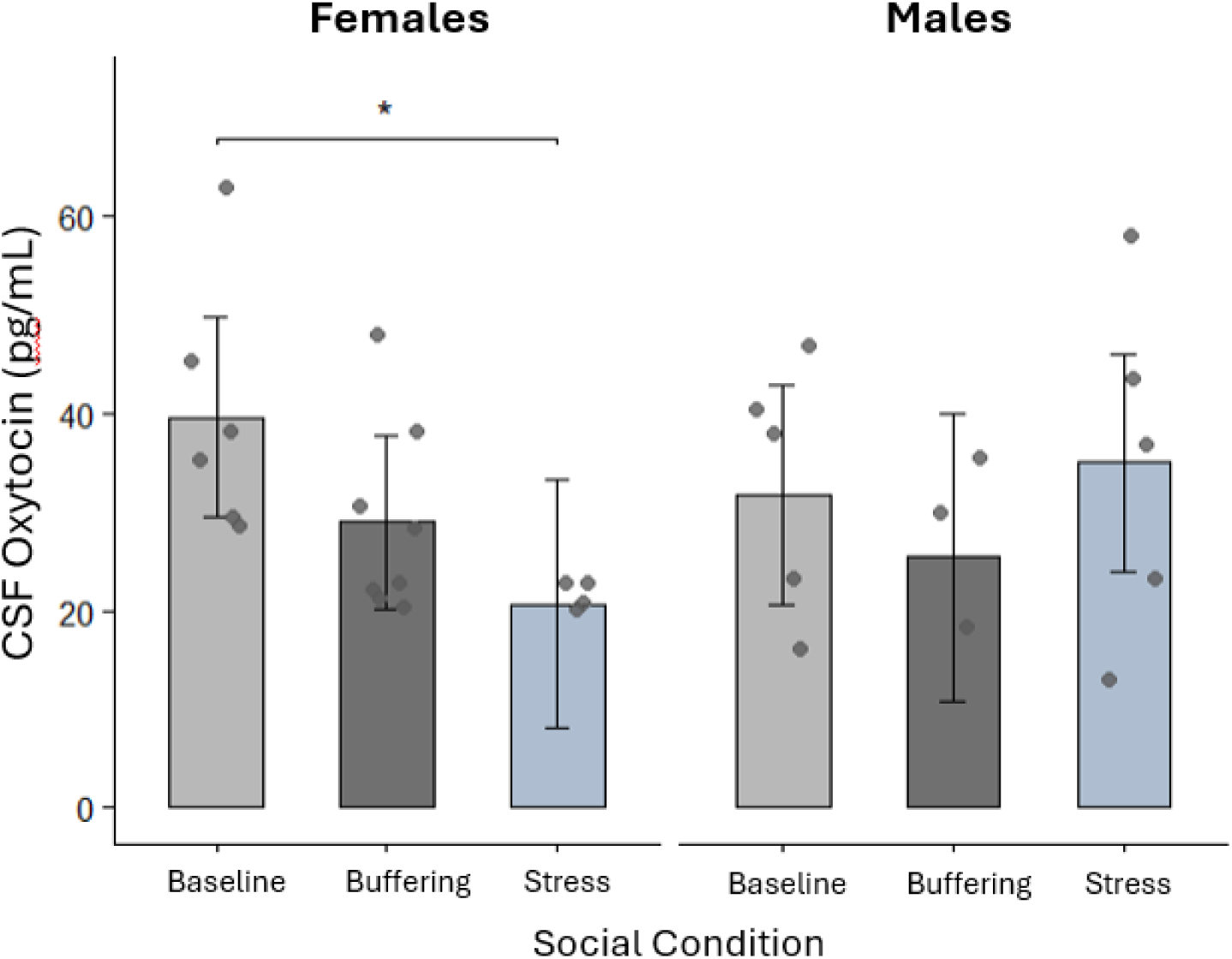
CSF OT Concentration by Sex and Condition. Mean CSF OT levels are shown for females and males under baseline, buffering, and stress conditions. The error bars represent standard error. Individual data points represent observed values for each subject. No significant main effects of condition (*F*(2, 17.23) = 1.79, *p* = .196) or sex (*F*(1, 13.29) = 0.005, *p* = .944) were observed. Planned comparisons indicated higher CSF OT levels in the baseline condition compared to stress in females (mean difference = 7.85, t = 1.07, **p = .049**\*, g = 0.62). * p<0.05

Type II ANOVA revealed no significant main effect of condition (F(2, 23.67) = 0.76, p = 0.480, partial η² = 0.06), no significant main effect of sex (F(1, 13.36) = 0.002, p = 0.966, partial η² = 0.001), and no significant interaction between condition and sex (F(2, 24.02) = 2.04, p = 0.152, partial η² = 0.15).

Exploratory comparisons showed that in females, BP_ND_ in the baseline condition was significantly higher than stress (estimate = 0.3450, SE = 0.167, df = 28.7, t = 2.066, **p = 0.048\***, g = 0.39). No other comparisons between experimental conditions revealed any significant differences.

#### Nucleus Accumbens (NAcc)

LMMs were used to examine the effects of condition, sex, and their interaction on NAcc BP_ND_. LMMs results are reported in table 2.

Type II ANOVA revealed no significant main effect of condition (F(2, 28.11) = 1.32, p = 0.281, partial η² = 0.07), no significant main effect of sex (F(1, 16.07) = 2.17, p = 0.159, partial η² = 0.12), and no significant condition × sex interaction (F(2, 28.65) = 0.404, p = 0.672, partial η² = 0.03).

Exploratory comparisons between conditions did not reveal any significant differences.

#### Septum

LMMs were used to examine the effects of condition, sex, and their interaction on septum BP_ND_. The septum model produced a singular fit, indicating that the random intercept term explained no additional variance beyond the fixed effects. Consequently, the conditional R² could not be calculated, and only the marginal R² value (0.088) is reported. LMMs results are reported in table 2.

Type II ANOVA revealed no significant main effect of condition (F(2, 44) = 0.6025, p = 0.5519, partial η² = 0.0267), no significant main effect of sex (F(1, 44) = 1.4893, p = 0.2288, partial η² = 0.0327), and no significant condition × sex interaction (F(2, 44) = 0.7056, p = 0.4993, partial η² = 0.0311).

Exploratory comparisons between conditions did not reveal any significant differences.

### 3.3 Cerebrospinal Fluid (CSF) Oxytocin (OT)

A LMM was fitted to examine the effects of condition and sex on CSF OT levels, with subject ID included as a random intercept. Random effects indicated moderate between-subject variability (σ² (ID) = 29.3, SD = 5.41), and residual variance was σ² = 111.0 (SD = 10.54).

A Type II ANOVA with Satterthwaite’s approximation for degrees of freedom revealed that the main effect of condition was not significant, (*F*(2, 17.23) = 1.79, *p* = .196, ηp² = 0.18) and the main effect of sex was also not significant, (*F*(1, 13.29) = 0.005, *p* = .944, ηp² = 0.002). The interaction between condition and sex approached significance, (*F*(2, 17.76) = 2.72, *p* = .093, ηp² = 0.24).

Tukey-adjusted pairwise comparisons indicated that, among females, CSF OT was significantly higher in the baseline condition compared to stress (mean difference = 7.85, *t* = 1.07, **p = .049\***, *g* = 0.62), reflecting a moderate effect size. No other significant contrasts were found.

### 3.4 Plasma Oxytocin (OT)

A LMM was fitted to examine the effects of condition and sex on plasma OT levels, with subject ID included as a random intercept. The model showed a singular fit, indicating negligible between-subject variability (σ² (ID) = 0.00), and residual variance was σ² = 101.3 (SD = 10.07). A Type III ANOVA with Satterthwaite’s approximation for degrees of freedom revealed that neither the main effect of condition (F(2, 45) = 0.89, p = .416, ηp² = 0.04), nor the main effect of sex (F(1, 45) = 0.19, p = .662, ηp² = 0.004), was significant. There were also no significant main effects of condition or sex and no significant interaction (F(2, 45) = 2.03, p = .14, ηp² = 0.08).

**Table 4:** Type III ANOVA results for plasma OT concentrations.

| Effect | Sum Sq | Mean Sq | NumDF | DenDF | F value | Pr(>F) | Partial $\eta^2$ |
| --- | --- | --- | --- | --- | --- | --- | --- |
| Condition | 181.22 | 90.611 | 2 | 45 | 0.894 | 0.416 | 0.04 |
| Sex | 19.62 | 19.619 | 1 | 45 | 0.194 | 0.662 | 0.004 |
| Condition x Sex | 411.93 | 205.96 | 2 | 45 | 0.326 | 0.14 | 0.08 |

**Figure 5:**
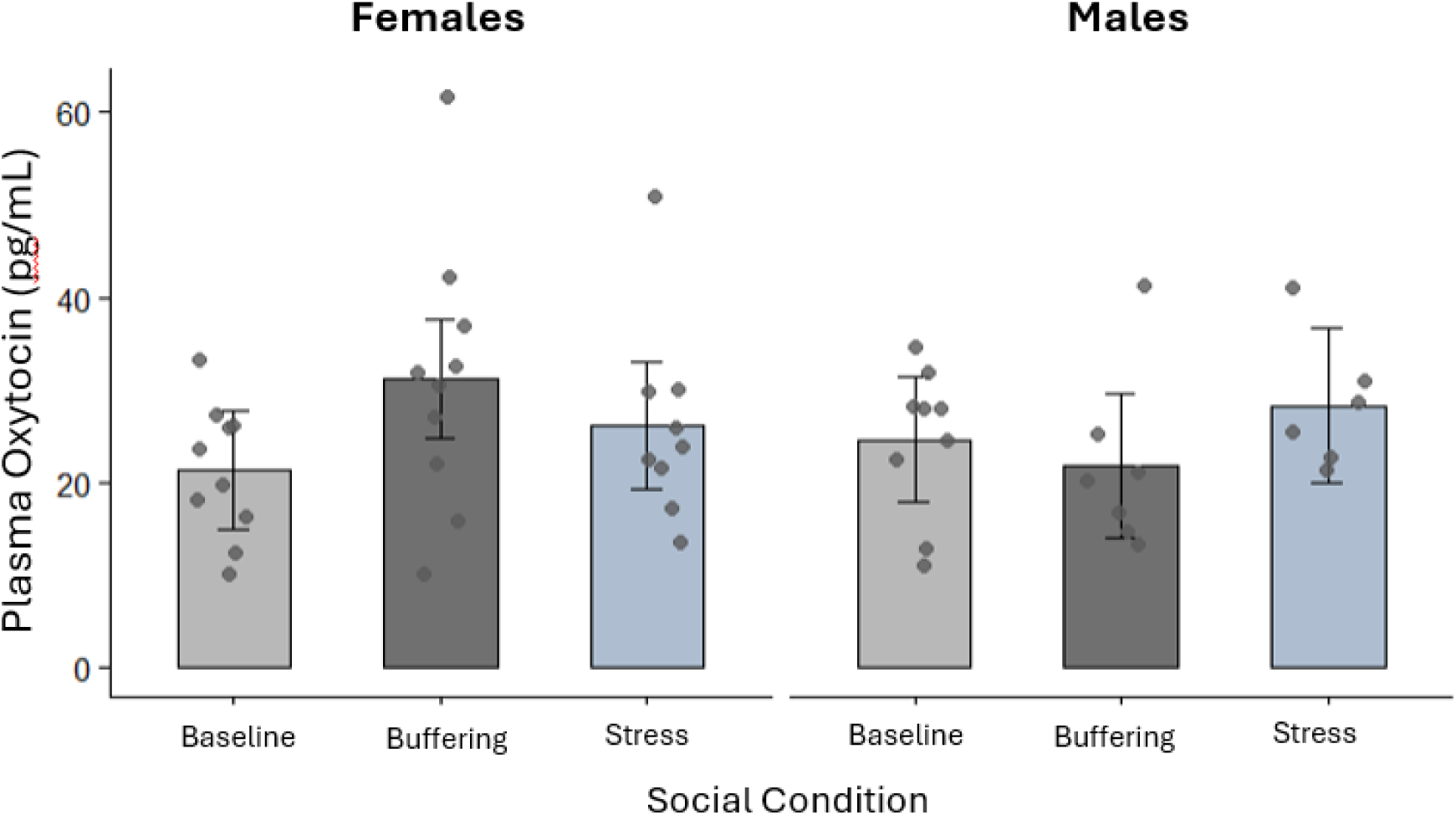
Plasma OT Concentrations by Condition and Sex. Mean plasma OT levels are shown for females and males under baseline, buffering, and stress conditions. The error bars represent standard error. Individual data points represent observed values for each subject. No significant main effects of condition (F(2, 45) = 0.89, p = .42) or sex (F(1, 45) = 0.19, p = .66) were found, and the condition × sex interaction was not significant, F(2, 45) = 2.03, p = .14.

### 3.5 Relationship between plasma and CSF OT

Spearman correlations between plasma and CSF OT levels revealed no significant associations overall (ρ = –0.32, *p* = .09) or within individual conditions. At baseline, plasma and CSF OT were weakly and positively correlated (ρ = 0.27, *p* = .45), whereas modest negative correlations were observed during the buffering (ρ = –0.56, *p* = .09) and stress (ρ = –0.36, *p* = .39) conditions. A linear model including condition as a moderator similarly revealed no significant plasma × condition interaction (F(5, 22) = 0.96, p = 0.47, R² = 0.18, adjusted R² = –0.01).

**Table 5:** Spearman’s rank correlations (ρ) between CSF and plasma OT levels during baseline, buffering, and stress conditions.

| Condition | $\rho$ (rho) | p |
| --- | --- | --- |
| Baseline | +0.27 | 0.446 |
| Buffering | -0.56 | 0.090 |
| Stress | -0.36 | 0.385 |

**Figure 6:**
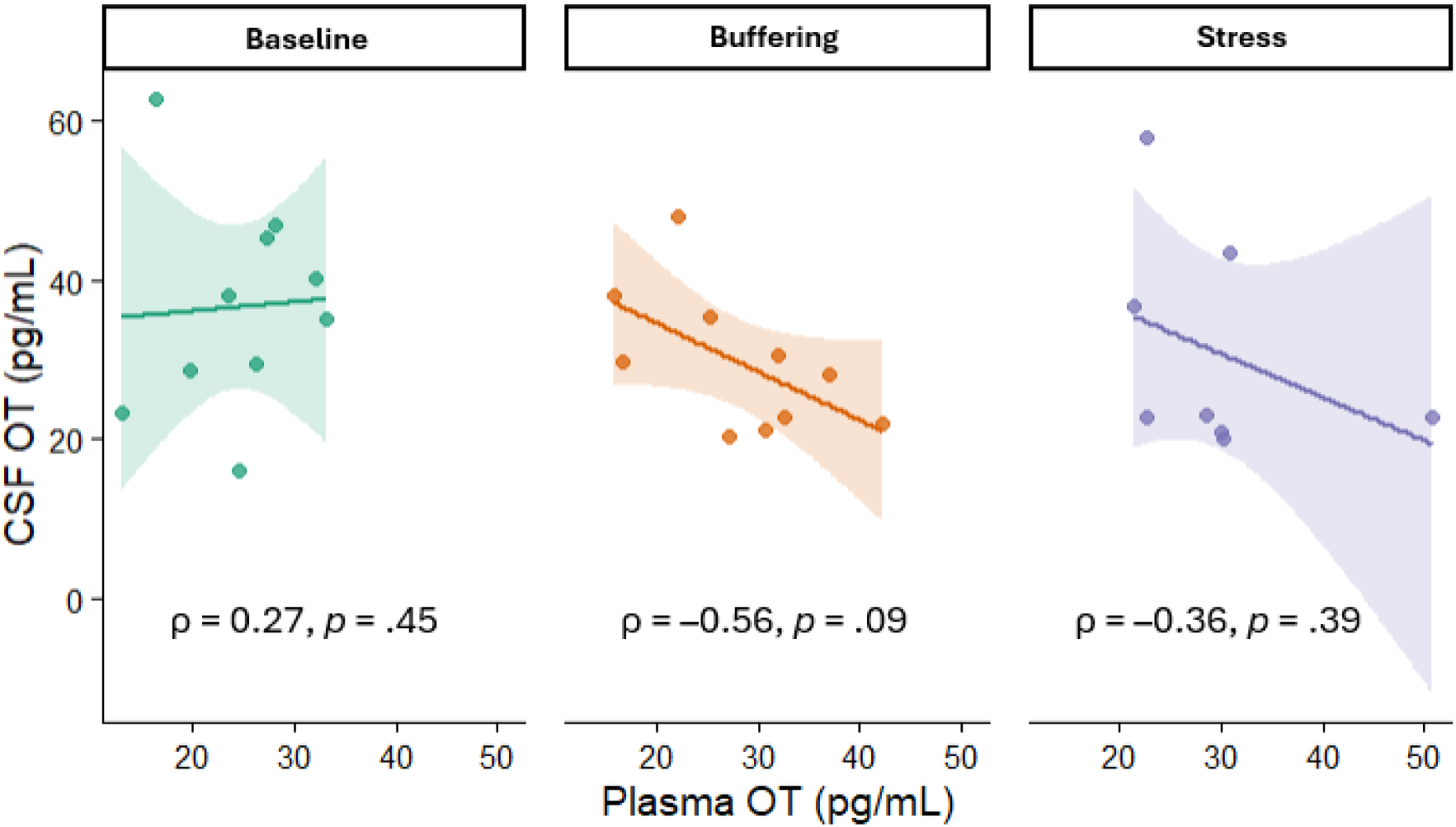
Associations between CSF and plasma OT across social conditions. Scatterplots show the associations between CSF and plasma OT during the baseline, buffering, and stress conditions. Lines represent Spearman correlations with shaded 95% confidence intervals. Correlation coefficients (ρ) and *p*-values are shown for each condition.

## Discussion

The findings from this study give us insight into the neurobiological mechanisms underlying social buffering in pair-bonded titi monkeys while undergoing an acute stressor. These findings provide overall support for a role of the κ-opioid system in stress response and social buffering, with important region- and sex-specific patterns.

The cortisol data support the prediction that separation from a mate leads to increased cortisol in both male and female titi monkeys (Mendoza et al., 2000). Exposure to an acute stressor (blood draw and short separation from pair mate) resulted in a significant increase in cortisol levels in the stress condition compared to baseline in both males and females. Moreover, no significant difference emerged between the baseline and buffering conditions, suggesting that the presence of a partner, after a stressor, maintained cortisol levels comparable to baseline. These findings align with previous studies demonstrating the stress-buffering effects of social support in both humans and animals and provide further evidence for the protective role of social relationships in stress regulation (Gobrogge & Wang, 2015; Lim & Hong, 2023; Smith & Wang, 2014).

PET imaging revealed condition- and sex-specific alterations in KOR availability across selected regions. In the amygdala, there was a trend toward a condition-by-sex interaction, with females exhibiting higher BP_ND_ of [¹¹C]GR103545 in the stress condition relative to baseline – a finding contrary to our hypothesis. We interpret the higher BP_ND_ as reflecting reduced endogenous dynorphin release, which would leave more KORs unoccupied and available for radiotracer binding, resulting in an increase in BP_ND_. In the amygdala, we expected higher release of endogenous dynorphin during the stress condition as stress typically induces dynorphin release (Bruchas et al., 2010; Knoll & Carlezon, 2010; Van’t Veer & Carlezon, 2013).

A potential explanation centers around the possibility that dynorphin – KOR signaling in the amygdala plays a more complex, circuit-dependent role than previously assumed. A recent study in mice demonstrated that KOR signaling within central nucleus of the amygdala (CeA) neurons is anxiolytic and facilitates accurate threat discrimination, rather than promoting stress responses (Baird et al., 2021). Importantly, the authors showed that these effects depend not on dynorphin produced locally within the CeA, but on dynorphin arriving from distal projections, primarily cortical and hypothalamic sources, which provide a tonic inhibitory influence through KOR activation. Their findings align with our data showing diminished dynorphin signaling under the stress condition relative to baseline. If stress disrupts the dynorphin input that normally reached the amygdala, fewer KORs would be activated endogenously, allowing our radioligand to bind to the receptors resulting in the higher BP_ND_ observed. Furthermore, we are interpreting higher BP_ND_ as a biomarker of reduced endogenous dynorphin release however alternative mechanisms such as changes in receptor density or affinity cannot be ruled out with the present data.

In the hippocampus, we found a significant main effect of sex, with follow-up comparisons indicating higher BP_ND_ in males than females specifically in the buffering condition. This pattern suggests that, in males, the presence of a mate may have been associated with reduced endogenous dynorphin release relative to females. The hippocampus plays a central role in stress regulation and in the neurobiology of pair bonds (Baxter et al., 2020; McEwen et al., 2015, 2016; Walum & Young, 2018). Thus, the observed effects may reflect a social buffering mechanism in which partner presence dampens dynorphin release during acute stress in males but not in females.

This sex-specific pattern in KOR binding fits within a broader literature documenting pronounced sex differences in κ-opioid pharmacology and pharmacokinetics (Rasakham & Liu-Chen, 2011; Russell et al., 2014). It is probable that variations in sex differences across species exist. In humans, women exhibit higher KOR availability compared to men, as evidence by a recent PET study investigating the relationship between the KOR system and social status (Matuskey et al., 2019). This finding aligns with other studies demonstrating that women have greater responses to KOR agonists than men (Gear et al., 1996, 1999). However, a PET study in humans found that men had higher KOR availability than women across most brain regions (Vijay et al., 2016). These differences may be attributed to the use of a KOR agonist vs. a KOR antagonist PET tracer (Johnson et al., 2023). In contrast, in rodents such as mice and rats, males tend to exhibit stronger effects of KOR activation on behavior compared to females (Russell et al., 2014). This is consistent with previous studies in rats and California mice reporting stronger effects of KOR activity on behavior in males than in females (Chartoff & Mavrikaki, 2015; Williams et al., 2018). Similarly, in rhesus monkeys, the κ-selective agonist U50,488 was more potent in males than in females, although sex differences in pharmacokinetics may have contributed to the different effects of U50,488 (Negus et al., 2002). Taken together, these findings show that sex differences in κ-opioid function are common across species, which aligns with the male-biased effect we observed in the hippocampus.

Another explanation may be sex-specific differences in stress responsiveness. In rodents, females generally exhibit a more pronounced neuroendocrine response to acute stress, showing higher cortisol and ACTH levels than males across a range of stressor types (Babb et al., 2013; Heck & Handa, 2019; Iwasaki-Sekino et al., 2009). In humans, many psychiatric disorders that are more prevalent in women are linked to stress, and these sex differences in stress responsiveness may contribute to the observed sex bias in disease (see Bangasser & Valentino, 2014). It is also possible that, in titi monkeys, females buffer their male partners more effectively after an acute stressor, resulting in reduced endogenous dynorphin release in the male hippocampus.

Contrary to our expectations, we did not find any significant effects of sex or condition in the septum and NAcc. Previous studies identified the lateral septum as being involved in the formation of pair bonds in titi monkeys and prairie voles (Bales et al., 2007; Hostetler et al., 2017; Y. Liu et al., 2001). A recent PET study in titi monkeys reported no significant reductions in KOR binding in the septum following administration of a κ-opioid antagonist (Almeida et al., 2025), suggesting that regions with moderate KOR binding may be less responsive to KOR blockade and therefore less likely to show measurable changes under the present imaging conditions.

The NAcc was of particular interest as it is strongly linked to both the OT (Dölen et al., 2013; Dölen & Malenka, 2014; Y. Liu & Wang, 2003) and opioidergic systems (Hakan & Henriksen, 1989; Henriksen & Willoch, 2008; Putnam & Chang, 2022; Trezza et al., 2011). In prairie voles, the long-term formation and maintenance of pair bond relies on both mu opioid receptors and KOR signaling (Resendez et al., 2012, 2013) in the NAcc shell where these receptors are thought to mediate reinforcing interactions with the preferred partner (MOR) and aversion towards strangers (KOR). Previous work has shown that activation of KORs in the NAcc shell promotes dysphoric behaviors (Pirino et al., 2020). A recent study in rats found that KOR activation in the caudal shell increases rearing and anxiety-like or avoidance behaviors, whereas activation in the rostral shell decreases anxiety-like behavior and promotes approach (Pirino et al., 2020). In this study, the lack of significant condition-related differences in NAcc KOR binding may be related to the functional organization of this region. Because the spatial resolution of the PET scans required the NAcc shell to be analyzed as a single region of interest in the present study, it was not possible to differentiate these subregions. Thus, the lack of observed differences in NAcc BP_ND_ does not necessarily rule out dynorphin involvement but may instead reflect subregion-specific effects that could not be examined with the present imaging approach.

In the hypothalamus, we found higher BP_ND_ in the baseline condition compared to the stress condition in females. Consistent with our hypothesis, we saw an increase of dynorphin release in the stress condition compared to baseline in females. Because the hypothalamus is the primary source of central OT release (Ludwig, 1998; Neumann, 2007), these imaging results provide important context for understanding the CSF OT and hypothalamic dynorphin patterns discussed in the following paragraph.

CSF OT levels did not show significant differences between conditions or sexes, although there was a trend towards a condition-sex interaction. In females, follow-up comparisons revealed CSF OT levels were higher in the baseline condition compared to the stress condition. As mentioned above, we also found a significant increase in hypothalamic dynorphin levels in females during the stress condition relative to baseline. These results align with the proposed mechanisms of separation distress outlined by Bales and Rogers (2022), who suggested that during acute separation, corticotropin-releasing hormone receptor type 2 (CRHR2) activation triggers dynorphin release and subsequent KOR activation, ultimately leading to suppression of OT release. Although the present study did not observe increased dynorphin levels in the NAcc shell as described in their model, the elevation of hypothalamic dynorphin in females during stress may reflect a related upstream process influencing oxytocinergic inhibition. Consistent with this interpretation, previous studies have shown that KOR activation can reduce OT release or signaling, supporting the idea that dynorphin-mediated KOR activation contributes to the stress-related downregulation of OT activity (Leng et al., 2008; Morris et al., 2010; Russell et al., 1993, 2003). Together, these results suggest that acute stress may transiently inhibit OT activity through a dynorphin–KOR pathway. The reason this effect was only observed in females remains speculative.

A possible explanation, aside from sex differences in κ-opioid function, lies in sex-specific differences in both neuroendocrine regulation and social behavior. While some studies have found central release of OT after a stressor (Babygirija et al., 2012; Nishioka et al., 1998), OT involvement may differ in acute versus chronic HPA activation as suggested by Bosch and colleagues (2016). They proposed that reducing OT activity during brief separations could serve as an adaptive function by motivating reunion with the partner, a mechanism that may have evolved to support maintenance of pair bonds.

Female and male titi monkeys contribute differently to the maintenance of the pair bond, with females contributing more than males to proximity and affiliation maintenance (Dolotovskaya et al., 2020). In titi monkeys, where males typically provide the majority of infant care and territorial protection, females may have a greater motivation to maintain the pair bond (Dolotovskaya et al., 2020), and may experience brief separations as more distressing, leading to stronger activation of the dynorphin-KOR system and subsequent inhibition of OT release. It should also be noted that some samples were insufficient for analysis, which may have reduced the robustness of our conclusions concerning CSF OT levels and the absence of effects observed in males.

Interestingly, the stress-related decrease in CSF OT in females was not accompanied by a corresponding change in plasma OT. This divergence is not unexpected, as central and peripheral OT are loosely coupled, and CSF is considered a closer indicator of central oxytocinergic activity. In addition, elevated concentrations of OT in the CSF are present for a longer period than in blood (Kendrick et al., 1991; Tabak et al., 2023), which may make central changes more detectable than peripheral ones. It is possible that the increase in hypothalamic dynorphin during the stress condition may reflect KOR activation acting directly on OT-producing neurons in the PVN or on their local terminals, reducing central OT release. Such effects would be expected to appear more clearly in CSF than in blood.

Finally, a central unresolved question in OT research concerns whether peripheral OT concentrations are predictive of central OT levels measured in CSF. Evidence from a meta-analysis of both human and animal studies indicates that central and peripheral OT levels are not associated at baseline, but it found a moderate positive association following stress in non-human primates (Valstad et al., 2017). Our current study found that plasma OT does not predict CSF OT, consistent with previous literature in humans (Kagerbauer et al., 2013) and macaques (Amico et al., 1990; Freeman et al., 2016). Thus, the evidence in titi monkeys suggests that plasma OT levels cannot be used as a reference for central OT levels measured in CSF, at least under our specific testing conditions.

Taken together, these findings suggest a model in which social buffering modulates κ-opioid receptor availability in a region- and sex-specific manner, rather than producing uniform changes in κ-opioid signaling across limbic regions. Consistent with prior social buffering literature, separation from a mate following a stressor was associated with elevated cortisol relative to baseline, whereas partner presence attenuated the endocrine stress response. In contrast, κ-opioid and OT responses varied across brain regions, experimental conditions, and sexes, highlighting the complexity of the neurobiological mechanisms underlying social buffering. Stress-related changes in OT were observed centrally but not peripherally, suggesting that plasma measures may be less sensitive to social context in titi monkeys.

Collectively, these results advance understanding of the neurobiology of stress regulation in pair-bonded species and provide the first evidence of sex differences in κ-opioid dynamics in titi monkeys, extending previous work that focused exclusively on males (Ragen et al., 2013, 2015b). At the same time, several limitations should be considered, including a modest sample size (N = 20) with substantial CSF sample loss (48%), which reduced statistical power for detecting sex-by-condition interactions; limited PET spatial resolution that precluded differentiation of functionally distinct brain subregions; potential effects of anesthesia on receptor binding and neurotransmitter dynamics; the inability to directly measure dynorphin release; and the use of post-hoc exploratory contrasts to probe region- and sex-specific effects, which increases the risk of Type I error. Despite these limitations, these findings contribute to our understanding of the protective role of social relationships in stress regulation across species, with implications for both animal and human studies.

## Author Contributions (CRediT)

*Claudia Manca* – investigation, project administration, visualization, formal analysis, data curation, writing – original draft.

*John P. Paulus* – investigation, visualization, formal analysis, writing – reviewing and editing.

*Alita J. D Almeida* – software, validation, writing – reviewing and editing.

*Anelise Caceres* – software, validation, writing – reviewing and editing.

*Meghan J. Sosnowski* – methodology, resources, supervision, writing – reviewing and editing.

*Brad A. Hobson* – software, validation, writing – reviewing and editing.

*Emilio Ferrer* – formal analysis, writing – reviewing and editing.

*Abhijit J. Chaudhari* – methodology, resources, supervision, funding acquisition, writing – reviewing and editing.

*Karen L. Bales* – conceptualization, methodology, resources, supervision, visualization, funding acquisition, writing – reviewing and editing.

## Acknowledgments

The authors would like to thank Joshua Waltenburg, Charles Smith and Sarah Tam for their assistance with PET scanning. The authors would also like to thank many members of the Bales lab who assisted with data collection; Gitanjali Gnanadesikan for validating the plasma oxytocin extraction procedure in titi monkeys and for developing the associated protocol; Jaleh Janatpour and Kevin Theis for animal care and the veterinary staff at CNPRC.

