## Supplementary material for "Brain Kappa Opioid Receptor Availability Across Stress and Social Buffering Conditions: A Positron Emission Tomography Study in Coppery Titi Monkeys": Suppl. Materials

**Supplementary Materials A**

2022-08-02_ArborOT_BasicAssayProcedure.docx

2022-08-02_RA-TitiMonkey-Protocol.docx

**Supplementary Materials B**

**Cortisol**

**Supplemental Table 1. Linear mixed model results for the effects of condition, sex, and their interaction on cortisol concentrations.**

| Predictors | Estimates | CI | p |
|---|---|---|---|
| (Intercept) | 626.60 | 452.19 – 801.02 | <0.001 |
| Buffering vs Baseline | 86.65 | -47.02 – 220.32 | 0.199 |
| Stress vs Baseline | 229.02 | 95.35 – 362.69 | **0.001*** |
| Male vs Female | -54.63 | -298.63 – 189.37 | 0.655 |
| Buffering × Male | 58.55 | -127.00 – 244.10 | 0.529 |
| Stress × Male | -53.34 | -238.89 – 132.21 | 0.566 |

Marginal $R^2$ = 0.103 / Conditional $R^2$ = 0.745

For the categorical predictor condition, the intercept corresponds to the baseline condition; for the categorical predictor sex, the intercept corresponds to females. Significant estimates and their *p*-values are bolded.

**CSF Oxytocin**

**Supplementary Table 2. Linear mixed model results for the effects of condition, sex, and their interaction on CSF OT concentrations.**

| Predictors | Estimates | CI | p |
|---|---|---|---|
| (Intercept) | 30.19 | 25.10 – 35.27 | <0.001 |
| Buffering vs Baseline | 5.43 | -0.35 – 11.22 | 0.064 |
| Stress vs Baseline | -3.04 | -9.27 – 3.19 | 0.323 |
| Male vs Female | -0.47 | -5.56 – 4.61 | 0.850 |
| Buffering × Male | 4.40 | -1.38 – 10.18 | 0.129 |
| Stress × Male | 2.26 | -3.97 – 8.49 | 0.460 |

Marginal $R^2$ = 0.103 / Conditional $R^2$ = 0.370

For the categorical predictor condition, the intercept corresponds to the baseline condition; for the categorical predictor sex, the intercept corresponds to females. Significant estimates and their *p*-values are bolded.

**Plasma Oxytocin**

**Supplementary Table 3. Linear mixed model results for the effects of condition, sex, and their interaction on plasma OT concentrations.**

| Predictors | Estimates | CI | p |
|---|---|---|---|
| (Intercept) | 25.56 | 22.66 – 28.45 | <0.001 |
| Buffering vs Baseline | -2.60 | -6.55 – 1.35 | 0.191 |
| Stress vs Baseline | 0.90 | -3.18 – 4.99 | 0.658 |
| Male vs Female | 0.63 | -2.26 – 3.52 | 0.662 |
| Buffering × Male | -2.33 | -6.27 – 1.62 | 0.241 |
| Stress × Male | 4.03 | -0.05 – 8.11 | 0.053 |

Marginal $R^2$ = 0.125 / Conditional $R^2$ = 0.028

For the categorical predictor condition, the intercept corresponds to the baseline condition; for the categorical predictor sex, the intercept corresponds to females. Significant estimates and their *p*-values are bolded.

**Brain Regions Tables and Results**

**Amygdala**

**Supplementary Table 4. Type III ANOVA Results for Amygdala BPnd**

| Effect | Sum Sq | Mean Sq | NumDF | DenDF | F Value | p-value | Partial $\eta^2$ |
|---|---|---|---|---|---|---|---|
| Condition | 0.104 | 0.052 | 2 | 25.982 | 1.890 | 0.171 | 0.127 |
| Sex | 0.017 | 0.017 | 1 | 15.633 | 0.632 | 0.439 | 0.038 |
| Condition x Sex | 0.180 | 0.090 | 2 | 25.982 | 3.250 | 0.055 | 0.200 |

**Supplementary Table 5. Pairwise Comparisons for Amygdala BPnd**

| Comparison | Estimate | SE | df | t-ratio | p-value | Hedges' g |
|---|---|---|---|---|---|---|
| Male Buffering vs Baseline | –0.15817 | 0.0817 | 26.7 | –1.936 | 0.0635 | –0.37449 |
| Male Buffering vs Stress | 0.13566 | 0.0775 | 27.2 | 1.749 | 0.0915 | 0.33528 |
| Male Baseline vs Stress | –0.02251 | 0.0809 | 27.7 | –0.278 | 0.7829 | –0.05291 |
| Female Buffering vs Baseline | –0.00915 | 0.0905 | 28.9 | –0.101 | 0.9202 | –0.01881 |
| Female Buffering vs Stress | –0.18346 | 0.1001 | 30.1 | –1.833 | 0.0767 | –0.33402 |
| Female Baseline vs Stress | –0.19260 | 0.0852 | 28.3 | –2.261 | **0.0316*** | –0.42519 |
| Buffering Males vs Females | –0.2265 | 0.118 | 38.4 | –1.912 | 0.0634 | –0.61687 |
| Baseline Males vs Females | –0.0775 | 0.109 | 34.4 | –0.710 | 0.4826 | –0.24221 |
| Stress Males vs Females | 0.0926 | 0.112 | 36.0 | 0.824 | 0.4152 | 0.27494 |
| Buffering vs Baseline (pooled) | –0.0837 | 0.0609 | 27.9 | –1.373 | 0.1808 | –0.25987 |
| Buffering vs Stress (pooled) | –0.0239 | 0.0633 | 29.0 | –0.377 | 0.7086 | –0.07010 |
| Baseline vs Stress (pooled) | –0.1076 | 0.0587 | 28.0 | –1.831 | 0.0777 | –0.34614 |

**Hippocampus**

**Supplementary Table 6. Type III ANOVA Results for Hippocampus BPnd**

| Effect | Sum Sq | Mean Sq | NumDF | DenDF | F Value | Raw p-value | Partial η² |
|---|---|---|---|---|---|---|---|
| Condition | 0.012409 | 0.006204 | 2 | 31.295 | 0.4106 | 0.66677 | 0.026 |
| Sex | 0.087198 | 0.087198 | 1 | 19.549 | 5.7707 | **0.02637*** | 0.23 |
| Condition x Sex | 0.045704 | 0.022852 | 2 | 31.295 | 1.5124 | 0.23606 | 0.088 |

**Supplementary Table 9. Pairwise Comparison for Hippocampus BPnd**

| Comparison | Estimate | SE | df | t-ratio | p-value | Hedges' g |
|---|---|---|---|---|---|---|
| Male Buffering vs Baseline | –0.10309 | 0.0602 | 27.4 | –1.711 | 0.0983 | –0.32686 |
| Male Buffering vs Stress | 0.10070 | 0.0570 | 27.8 | 1.767 | 0.0882 | 0.33539 |
| Male Baseline vs Stress | –0.00239 | 0.0593 | 28.5 | –0.040 | 0.9681 | –0.00756 |
| Female Buffering vs Baseline | 0.02348 | 0.0659 | 30.6 | 0.356 | 0.7240 | 0.06444 |
| Female Buffering vs Stress | –0.04644 | 0.0724 | 32.7 | –0.641 | 0.5257 | –0.11217 |
| Female Baseline vs Stress | –0.02296 | 0.0622 | 29.5 | –0.369 | 0.7148 | –0.06789 |
| Baseline Males vs Females | –0.0816 | 0.0676 | 41.0 | –1.206 | 0.2346 | –0.37665 |
| Buffering Males vs Females | –0.2082 | 0.0750 | 42.8 | –2.776 | **0.0081*** | –0.84917 |
| Stress Males vs Females | –0.0610 | 0.0702 | 41.8 | –0.869 | 0.3896 | –0.26898 |
| Buffering vs Baseline (pooled) | –0.0398 | 0.0446 | 29.1 | –0.892 | 0.3798 | –0.16531 |
| Buffering vs Stress (pooled) | 0.0271 | 0.0461 | 30.7 | 0.589 | 0.5602 | 0.10622 |
| Baseline vs Stress (pooled) | –0.0127 | 0.0430 | 29.1 | –0.295 | 0.7702 | –0.05471 |

**Hypothalamus**

**Supplementary Table 10. Type II ANOVA Results for Hypothalamus BPnd**

| Effect | Sum Sq | Mean Sq | NumDF | DenDF | F value | Raw p-value | Partial η² |
|---|---|---|---|---|---|---|---|
| Condition | 0.16227 | 0.081136 | 2 | 23.674 | 0.7581 | 0.4796 | 0.06 |
| Sex | 0.0002 | 0.0002 | 1 | 13.264 | 0.0019 | 0.9662 | 0.0001 |
| Condition x Sex | 0.4368 | 0.2184 | 2 | 24.041 | 2.0405 | 0.1519 | 0.15 |

**Supplementary Table 11. Pairwise Comparison for Hypothalamus BPnd**

| Comparison | Estimate | SE | df | t-ratio | Raw p-value | Hedges' g |
|---|---|---|---|---|---|---|
| Male Buffering vs Baseline | –0.1766 | 0.161 | 26.9 | –1.100 | 0.2812 | –0.21186 |
| Male Buffering vs Stress | 0.0665 | 0.152 | 27.4 | 0.437 | 0.6658 | 0.08340 |
| Male Baseline vs Stress | –0.1101 | 0.159 | 27.9 | –0.694 | 0.4937 | –0.13118 |
| Female Buffering vs Baseline | 0.1131 | 0.177 | 29.5 | 0.638 | 0.5282 | 0.11761 |

| | | | | | | |
|---|---|---|---|---|---|---|
| Female Buffering vs Stress | 0.2319 | 0.196 | 31.0 | 1.185 | 0.2449 | 0.21295 |
| Female Baseline vs Stress | 0.3450 | 0.167 | 28.7 | 2.066 | **0.0479*** | 0.38561 |
| Baseline Males vs Females | 0.2331 | 0.200 | 37.0 | 1.164 | 0.2518 | 0.38278 |
| Buffering Males vs Females | –0.0566 | 0.219 | 40.4 | –0.258 | 0.7976 | –0.08125 |
| Stress Males vs Females | –0.2220 | 0.207 | 38.4 | –1.073 | 0.2898 | –0.34651 |

**NAcc**

**Supplementary Table 12. Type II ANOVA Results for NAcc BPnd**

| Effect | Sum Sq | Mean Sq | NumDF | DenDF | F Value | Raw p-value | Partial η² |
|---|---|---|---|---|---|---|---|
| Condition | 0.11936 | 0.05967 | 2 | 28.113 | 1.3249 | 0.2819 | 0.07 |
| Sex | 0.09819 | 0.09819 | 1 | 16.007 | 2.1798 | 0.1592 | 0.12 |
| Condition x Sex | 0.03637 | 0.01819 | 2 | 28.650 | 0.4037 | 0.6716 | 0.03 |

**Supplementary Table 13. Pairwise Comparison for NAcc BPnd**

| Comparison | Estimate | SE | df | t-ratio | Raw p-value | Hedges' g |
|---|---|---|---|---|---|---|
| Male Buffering vs Baseline | –0.12589 | 0.1039 | 27.6 | –1.211 | 0.2361 | –0.23055 |
| Male Buffering vs Stress | 0.13880 | 0.0983 | 27.9 | 1.413 | 0.1688 | 0.26755 |
| Male Baseline vs Stress | 0.01291 | 0.1022 | 28.8 | 0.126 | 0.9003 | 0.02357 |
| Female Buffering vs Baseline | –0.10209 | 0.1133 | 31.0 | –0.901 | 0.3744 | –0.16189 |
| Female Buffering vs Stress | 0.00945 | 0.1242 | 33.3 | 0.076 | 0.9398 | 0.01318 |
| Female Baseline vs Stress | –0.09264 | 0.1071 | 29.8 | –0.865 | 0.3939 | –0.15836 |
| Baseline Males vs Females | –0.1449 | 0.113 | 42.1 | –1.281 | 0.2072 | –0.39493 |
| Buffering Males vs Females | –0.1687 | 0.126 | 43.2 | –1.341 | 0.1869 | –0.40785 |
| Stress Males vs Females | –0.0394 | 0.118 | 42.6 | –0.335 | 0.7394 | –0.10257 |
| Pooled: Baseline vs Buffering | –0.1140 | 0.0769 | 29.4 | –1.483 | 0.1487 | –0.27348 |
| Pooled: Baseline vs Stress | –0.0399 | 0.0740 | 29.3 | –0.539 | 0.5942 | –0.09947 |
| Pooled: Buffering vs Stress | 0.0741 | 0.0792 | 31.1 | 0.936 | 0.3564 | 0.16773 |

**Septum**

**Supplementary Table 14. Type II ANOVA Results for Septum BPnd**

| Effect | Sum Sq | Mean Sq | NumDF | DenDF | F Value | Raw p-value | Partial η² |
|---|---|---|---|---|---|---|---|
| Condition | 0.09850 | 0.04925 | 2 | 44 | 0.6257 | 0.540 | 0.028 |
| Sex | 0.12522 | 0.12521 | 1 | 44 | 1.5908 | 0.214 | 0.035 |

| | | | | | | | |
|---|---|---|---|---|---|---|---|
| Condition x Sex | 0.11107 | 0.05553 | 2 | 44 | 0.7056 | 0.499 | 0.031 |

**Supplementary Table 15. Pairwise Comparison for Septum BPnd**

| Comparison | Estimate | SE | df | t-ratio | Raw p-value | Hedges' g |
|---|---|---|---|---|---|---|
| Male Buffering vs Baseline | –0.0415 | 0.137 | 28.6 | –0.304 | 0.7636 | –0.05679 |
| Male Buffering vs Stress | 0.0688 | 0.129 | 28.4 | 0.533 | 0.5983 | 0.09988 |
| Male Baseline vs Stress | 0.0272 | 0.134 | 29.7 | 0.204 | 0.8399 | 0.03739 |
| Female Buffering vs Baseline | –0.1517 | 0.147 | 32.8 | –1.035 | 0.3084 | –0.18072 |
| Female Buffering vs Stress | –0.0490 | 0.159 | 35.8 | –0.309 | 0.7592 | –0.05166 |
| Female Baseline vs Stress | –0.2007 | 0.139 | 31.1 | –1.440 | 0.1598 | –0.25808 |
| Baseline Males vs Females | –0.2112 | 0.134 | 44 | –1.580 | 0.1213 | –0.47634 |
| Buffering Males vs Females | –0.1011 | 0.150 | 44 | –0.675 | 0.5029 | –0.20364 |
| Stress Males vs Females | 0.0167 | 0.139 | 44 | 0.120 | 0.9051 | 0.03616 |
| Pooled: Baseline vs Buffering | –0.09660 | 0.1002 | 30.8 | –0.964 | 0.3427 | –0.17368 |
| Pooled: Baseline vs Stress | –0.08673 | 0.0966 | 30.4 | –0.898 | 0.3762 | –0.16276 |
| Pooled: Buffering vs Stress | 0.00987 | 0.1023 | 32.8 | 0.096 | 0.9238 | 0.01684 |

### Exploratory Brain Regions Results and Tables

#### Cingulate

The buffering condition showed a significant decrease in BPnd relative to baseline ($\beta = –0.1036$, SE = 0.0405, t = –2.561, **p = 0.0162***), and the stress condition showed a significant increase relative to baseline ($\beta = 0.1139$, SE = 0.0435, t = 2.619, **p = 0.0139***). The main effect of sex was not significant ($\beta = –0.0354$, SE = 0.0535, t = –0.661, p = 0.517), and neither interaction term reached significance. Model $R^2$ values were $R^2$ marginal = 0.121 and $R^2$ conditional = 0.558.

Type III ANOVA revealed a significant main effect of condition (F(2, 28.60) = 4.47, **p = 0.020***, FDR-corrected p = 0.238, partial $\eta^2 = 0.123$). The main effect of sex was not significant (FDR-corrected p = 0.763, partial $\eta^2 = 0.023$), and the condition × sex interaction was also not significant (FDR-corrected p = 0.816, partial $\eta^2 = 0.023$).

Post-hoc pairwise comparisons showed that, after FDR correction, no contrast remained statistically significant. Significant pairwise contrasts prior to FDR correction are reported in Supplementary Table 18.

**Supplementary Table 16. Linear Mixed Model (LMM) of Cingulate BPnd**

| Predictor | Estimate (β) | Std. Error (SE) | t(df) | t value | p-value |
|---|---|---|---|---|---|

| | | | | | |
|---|---|---|---|---|---|
| Intercept | 1.69846 | 0.05347 | 18.29 | 31.763 | < 0.001 |
| Condition (Buffering) | –0.10364 | 0.04048 | 27.91 | –2.561 | **0.0162*** |
| Condition (Stress) | 0.11393 | 0.04350 | 28.94 | 2.619 | **0.0139*** |
| Sex (Male) | –0.03535 | 0.05347 | 18.29 | –0.661 | 0.5168 |
| Buffering × Sex (Male) | 0.01653 | 0.04048 | 27.91 | 0.408 | 0.6861 |
| Stress × Sex (Male) | –0.03605 | 0.04350 | 28.94 | –0.829 | 0.4139 |

**Supplementary Table 17. Type III ANOVA Results for Cingulate BPnd**

| Effect | Sum Sq | Mean Sq | NumDF | DenDF | F | p (raw) | p (FDR) | partial η² |
|---|---|---|---|---|---|---|---|---|
| Condition | 0.356 | 0.178 | 2 | 28.6 | 4.471 | **0.020*** | 0.238 | 0.123 |
| Sex | 0.017 | 0.017 | 1 | 18.2 | 0.437 | 0.517 | 0.763 | 0.023 |
| Condition x Sex | 0.027 | 0.014 | 2 | 28.6 | 0.344 | 0.712 | 0.816 | 0.023 |

**Supplementary Table 18. Pairwise Comparisons for Cingulate BPnd**

| Comparison | Estimate | SE | df | t-ratio | Raw p-value | FDR p-value | Hedges' g |
|---|---|---|---|---|---|---|---|
| Female Buffering vs Baseline | −0.165 | 0.109 | 28.9 | −1.520 | 0.139 | 0.418 | −0.283 |
| Female Buffering vs Stress | 0.069 | 0.120 | 30.1 | 0.572 | 0.572 | 0.763 | 0.104 |
| Female Baseline vs Stress | −0.096 | 0.102 | 28.3 | −0.943 | 0.354 | 0.617 | −0.177 |
| Male Buffering vs Baseline | −0.270 | 0.098 | 26.7 | −2.757 | **0.010*** | 0.123 | −0.533 |
| Male Buffering vs Stress | 0.180 | 0.093 | 27.2 | 1.933 | 0.064 | 0.255 | 0.370 |
| Male Baseline vs Stress | −0.090 | 0.097 | 27.6 | −0.931 | 0.360 | 0.617 | −0.177 |
| Baseline Males vs Females | −0.038 | 0.131 | 34.2 | −0.286 | 0.776 | 0.816 | −0.098 |
| Buffering Males vs Females | −0.143 | 0.143 | 38.3 | −1.001 | 0.323 | 0.617 | −0.324 |
| Stress Males vs Females | −0.032 | 0.135 | 35.8 | −0.234 | 0.816 | 0.816 | −0.078 |

| | | | | | | | |
|---|---|---|---|---|---|---|---|
| Pooled: Baseline vs Buffering | −0.218 | 0.073 | 27.9 | −2.976 | **0.006*** | 0.078 | −0.563 |
| Pooled: Baseline vs Stress | −0.093 | 0.071 | 28.0 | −1.325 | 0.196 | 0.420 | −0.250 |
| Pooled: Buffering vs Stress | 0.124 | 0.076 | 29.0 | 1.635 | 0.113 | 0.338 | 0.304 |

**Claustrum**

The buffering condition showed a marginal decrease relative to baseline (β = –0.0923, SE = 0.0515, t = –1.791, p = 0.0835), whereas the stress condition showed a significant increase relative to baseline (β = 0.1336, SE = 0.0549, t = 2.433, **p = 0.0209***). There was also a significant main effect of sex, with males showing lower BPnd than females (β = –0.1202, SE = 0.0509, t = –2.359, **p = 0.0293***). Neither interaction term was significant. Model $R^2$ values were $R^2$ marginal = 0.248 and $R^2$ conditional = 0.446.
Type III ANOVA revealed a marginal main effect of condition (F(2, 30.55) = 3.17, p = 0.056, partial $\eta^2$ = 0.172) and a significant main effect of sex before correction (F(1, 18.71) = 5.57, **p = 0.029***, partial $\eta^2$ = 0.229). However, none of the ANOVA effects remained significant after FDR correction (corrected ps ≥ 0.131). The condition × sex interaction was not significant (corrected p = 0.573).
Post-hoc pairwise comparisons showed that, after FDR correction, no contrast remained statistically significant. Significant pairwise contrasts prior to FDR correction are reported in Supplementary Table 21.

**Supplementary Table 19. Linear Mixed Model (LMM) of Claustrum BPnd**

| Fixed Effect | Estimate | Std. Error | t(df) | t value | p-value |
|---|---|---|---|---|---|
| Intercept | 2.06850 | 0.05094 | 18.71 | 40.603 | < 0.001 |
| Condition (Buffering) | –0.09225 | 0.05150 | 29.50 | –1.791 | 0.0835 |
| Condition (Stress) | 0.13362 | 0.05493 | 31.16 | 2.433 | **0.0209*** |
| Sex (Male) | –0.12019 | 0.05094 | 18.71 | –2.359 | **0.0293*** |
| Buffering × Sex (Male) | –0.01506 | 0.05150 | 29.50 | –0.293 | 0.7719 |
| Stress × Sex (Male) | –0.05025 | 0.05493 | 31.16 | –0.915 | 0.3673 |

**Supplementary Table 20. Type III ANOVA Results for Claustrum BPnd**

| Effect | Sum Sq | Mean Sq | NumDF | DenDF | F | p (raw) | p (FDR) | partial $\eta^2$ |
|---|---|---|---|---|---|---|---|---|
| Condition | 0.415 | 0.208 | 2 | 30.548 | 3.172 | 0.056 | 0.134 | 0.172 |
| Sex | 0.365 | 0.365 | 1 | 18.708 | 5.566 | **0.029*** | 0.131 | 0.229 |
| Condition x Sex | 0.105 | 0.052 | 2 | 30.548 | 0.799 | 0.459 | 0.573 | 0.050 |

**Supplementary Table 21. Pairwise Comparisons for Claustrum BPnd**

| Comparison | Estimate | SE | df | t-ratio | Raw p-value | FDR p-value | Hedges' g |
|---|---|---|---|---|---|---|---|
| Female Buffering vs Baseline | −0.191 | 0.137 | 30.6 | −1.390 | 0.175 | 0.291 | −0.251 |
| Female Buffering vs Stress | 0.059 | 0.151 | 32.6 | 0.394 | 0.696 | 0.746 | 0.069 |
| Female Baseline vs Stress | −0.131 | 0.130 | 29.5 | −1.013 | 0.319 | 0.479 | −0.186 |
| Male Buffering vs Baseline | −0.261 | 0.125 | 27.4 | −2.081 | **0.047*** | 0.134 | −0.398 |
| Male Buffering vs Stress | 0.291 | 0.119 | 27.7 | 2.449 | **0.021*** | 0.131 | 0.465 |
| Male Baseline vs Stress | 0.030 | 0.124 | 28.5 | 0.239 | 0.813 | 0.813 | 0.045 |
| Baseline Males vs Females | −0.271 | 0.141 | 40.9 | −1.915 | 0.062 | 0.134 | −0.599 |
| Buffering Males vs Females | −0.341 | 0.157 | 42.7 | −2.178 | **0.035*** | 0.131 | −0.667 |
| Stress Males vs Females | −0.110 | 0.147 | 41.7 | −0.749 | 0.458 | 0.573 | −0.232 |
| Pooled: Baseline vs Buffering | −0.226 | 0.093 | 29.1 | −2.430 | **0.022*** | 0.131 | −0.451 |
| Pooled: Baseline vs Stress | −0.051 | 0.090 | 29.0 | −0.568 | 0.574 | 0.663 | −0.105 |
| Pooled: Buffering vs Stress | 0.175 | 0.096 | 30.7 | 1.824 | 0.078 | 0.146 | 0.329 |

### Orbitofrontal Cortex

The buffering condition showed a significant decrease relative to baseline ($\beta$ = –0.125, SE = 0.054, t = –2.309, **p = 0.028***), and the stress condition showed a significant increase relative to baseline ($\beta$ = 0.134, SE = 0.058, t = 2.334, **p = 0.026***). The main effect of sex was not significant ($\beta$ = –0.042, SE = 0.045, t = –0.923, p = 0.368), and neither interaction term reached significance (buffering × sex: $\beta$ = –0.002, SE = 0.054, t = –0.034, p = 0.974; stress × sex: $\beta$ = –0.025, SE = 0.058, t = –0.433, p = 0.668). Model $R^2$ values were $R^2$ marginal = 0.152 and $R^2$ conditional = 0.252.

Type III ANOVA revealed a significant main effect of condition before correction (F(2, 31.94) = 3.576, **p = 0.040**, partial $\eta^2$ = 0.183), but the corrected p-value did not reach significance (FDR-corrected p = 0.210). The main effect of sex was not significant (F(1, 18.99) = 0.852, p = 0.368, partial $\eta^2$ = 0.043; FDR-corrected p = 0.584), and the interaction between condition and sex was also not significant (F(2, 31.94) = 0.137, p = 0.873, partial $\eta^2$ = 0.008; FDR-corrected p = 0.873).

Post-hoc pairwise comparisons showed that, after FDR correction, no contrast remained statistically significant. Significant pairwise contrasts prior to FDR correction are reported in Supplementary Table 24.

**Supplementary Table 22. Linear Mixed Model (LMM) of Orbitofrontal Cortex BPnd**

| Fixed Effect | Estimate | Std. Error | df | t value | p-value |
|---|---|---|---|---|---|
| Intercept | 1.673 | 0.082 | 38.12 | 20.295 | <0.001 |
| Condition (Buffering) | 0.061 | 0.127 | 30.16 | 0.479 | 0.636 |
| Condition (Stress) | 0.152 | 0.112 | 28.02 | 1.355 | 0.186 |
| Sex (Male) | 0.079 | 0.123 | 39.39 | 0.641 | 0.526 |
| Buffering × Sex (Male) | 0.193 | 0.170 | 28.69 | 1.134 | 0.266 |
| Stress × Sex (Male) | -0.054 | 0.155 | 27.54 | -0.350 | 0.729 |

**Supplementary Table 23. Type III ANOVA Results for Orbitofrontal Cortex BPnd**

| Effect | Sum Sq | Mean Sq | NumDF | DenDF | F | p (raw) | p (FDR) | partial $\eta^2$ |
|---|---|---|---|---|---|---|---|---|
| Condition | 0.526 | 0.263 | 2 | 31.937 | 3.576 | **0.040*** | 0.210 | 0.183 |
| Sex | 0.063 | 0.063 | 1 | 18.993 | 0.852 | 0.368 | 0.584 | 0.043 |
| Condition × Sex | 0.020 | 0.010 | 2 | 31.937 | 0.137 | 0.873 | 0.873 | 0.008 |

**Supplementary Table 24. Pairwise Comparisons for Orbitofrontal Cortex BPnd**

| Comparison | Estimate | SE | df | t-ratio | Raw p-value | FDR p-value | Hedges' g |
|---|---|---|---|---|---|---|---|
| Female Buffering vs Baseline | −0.236 | 0.144 | 31.7 | −1.646 | 0.110 | 0.394 | −0.292 |

| | | | | | | | |
|---|---|---|---|---|---|---|---|
| Female Buffering vs Stress | 0.092 | 0.157 | 34.4 | 0.584 | 0.563 | 0.649 | 0.100 |
| Female Baseline vs Stress | −0.145 | 0.136 | 30.4 | −1.064 | 0.296 | 0.554 | −0.193 |
| Male Buffering vs Baseline | −0.283 | 0.133 | 28.0 | −2.130 | **0.042*** | 0.210 | −0.403 |
| Male Buffering vs Stress | 0.195 | 0.125 | 28.1 | 1.555 | 0.131 | 0.394 | 0.293 |
| Male Baseline vs Stress | −0.088 | 0.130 | 29.1 | −0.674 | 0.505 | 0.649 | −0.125 |
| Baseline Males vs Females | −0.087 | 0.137 | 43.4 | −0.635 | 0.529 | 0.649 | −0.193 |
| Buffering Males vs Females | −0.133 | 0.153 | 43.8 | −0.869 | 0.390 | 0.584 | −0.263 |
| Stress Males vs Females | −0.030 | 0.143 | 43.6 | −0.211 | 0.834 | 0.873 | −0.064 |
| Pooled: Baseline vs Buffering | −0.259 | 0.098 | 30.0 | −2.654 | **0.013*** | 0.189 | −0.485 |
| Pooled: Baseline vs Stress | −0.116 | 0.094 | 29.8 | −1.235 | 0.226 | 0.485 | −0.226 |
| Pooled: Buffering vs Stress | 0.143 | 0.100 | 31.9 | 1.427 | 0.163 | 0.408 | 0.253 |

**Thalamus**

None of the fixed effects reached significance at the model level. The buffering condition did not significantly differ from baseline ($\beta = -0.027$, SE = 0.016, $t = -1.672$, $p = 0.105$), and the stress condition also did not differ from baseline ($\beta = 0.014$, SE = 0.017, $t = 0.805$, $p = 0.427$). Sex was not a significant predictor ($\beta = -0.007$, SE = 0.013, $t = -0.554$, $p = 0.586$), and neither interaction term reached significance (buffering × sex: $\beta = -0.005$, SE = 0.016, $t = -0.325$, $p = 0.748$; stress × sex: $\beta = -0.012$, SE = 0.017, $t = -0.701$, $p = 0.489$). Model $R^2$ values were $R^2$ marginal = 0.081 and $R^2$ conditional = 0.165.

Type II ANOVA indicated no significant main effects or interactions. The main effect of condition was not significant ($F(2, 30.18) = 1.434$, $p = 0.254$, partial $\eta^2 = 0.087$; FDR-corrected $p = 0.696$). Sex was also not significant ($F(1, 17.21) = 0.279$, $p = 0.604$, partial $\eta^2 = 0.016$; FDR-corrected $p = 0.696$). The condition × sex interaction did not reach significance ($F(2, 30.80) = 0.557$, $p = 0.579$, partial $\eta^2 = 0.035$; FDR-corrected $p = 0.696$). As there were no significant main effects or interactions, pairwise comparisons are not reported.

**Supplementary Table 25. Linear Mixed Model (LMM) of Thalamus BPnd**

| Predictor | Estimate | Std. Error | df | t value | p-value |
|---|---|---|---|---|---|
| Intercept | 0.402 | 0.022 | 42 | 18.237 | <0.001 |
| Condition (Buffering) | 0.031 | 0.036 | 42 | 0.864 | 0.392 |
| Condition (Stress) | 0.027 | 0.036 | 42 | 0.757 | 0.453 |
| Sex (Male) | 0.024 | 0.033 | 42 | 0.735 | 0.466 |
| Buffering × Sex (Male) | -0.013 | 0.050 | 42 | -0.251 | 0.803 |
| Stress × Sex (Male) | -0.011 | 0.049 | 42 | -0.228 | 0.821 |

**Supplementary Table 26. Type II ANOVA Results for Thalamus BPnd**

| Effect | Sum Sq | Mean Sq | NumDF | DenDF | F | p (raw) | p (FDR) | partial η² |
|---|---|---|---|---|---|---|---|---|
| Condition | 0.01810 | 0.00905 | 2 | 30.18 | 1.434 | 0.254 | 0.696 | 0.087 |
| Sex | 0.00176 | 0.00176 | 1 | 17.21 | 0.279 | 0.604 | 0.696 | 0.016 |
| Condition × Sex | 0.00703 | 0.00351 | 2 | 30.80 | 0.557 | 0.579 | 0.696 | 0.035 |

**Putamen**

The buffering condition did not significantly differ from baseline (β = –0.042, SE = 0.033, t = –1.26, p = 0.219), whereas the stress condition showed a significant increase relative to baseline (β = 0.082, SE = 0.035, t = 2.33, **p = 0.026*).** Sex was also a significant predictor, with males exhibiting lower BPnd than females overall (β = –0.075, SE = 0.035, t = –2.13, **p = 0.046***). Neither the buffering × sex interaction (p = 0.932) nor the stress × sex interaction (p = 0.248) reached significance. Model $R^2$ values were $R^2$ marginal = 0.221 and $R^2$ conditional = 0.478.

Type III ANOVA showed no main effect of condition (F(2, 30.80) = 2.72, p = 0.082, FDR-corrected p = 0.175, partial $\eta^2$ = 0.150) and a significant main effect of sex (F(1, 19.55) = 4.54, p = 0.046, FDR-corrected p = 0.138, partial $\eta^2$ = 0.189). The condition × sex interaction was not significant (F(2, 30.80) = 1.00, p = 0.380, FDR-corrected p = 0.570, partial $\eta^2$ = 0.061).

Post-hoc pairwise comparisons showed that, after FDR correction, no contrast remained statistically significant. Significant pairwise contrasts prior to FDR correction are reported in Supplementary Table 29.

**Supplementary Table 27. Linear Mixed Model (LMM) of Putamen BPnd**

| Predictor | Estimate | Std. Error | df | t value | p-value |
|---|---|---|---|---|---|
| Intercept | 1.436 | 0.035 | 19.545 | 40.559 | < 0.001 |
| Buffering vs Baseline | -0.042 | 0.033 | 29.874 | -1.256 | 0.219 |

| | | | | | |
|---|---|---|---|---|---|
| Stress vs Baseline | 0.082 | 0.035 | 31.309 | 2.330 | **0.026*** |
| Sex (Male) | -0.075 | 0.035 | 19.545 | -2.132 | **0.046*** |
| Buffering × Male | -0.003 | 0.033 | 29.874 | -0.086 | 0.932 |
| Stress × Male | -0.042 | 0.035 | 31.309 | -1.176 | 0.248 |

**Supplementary Table 28. Type III ANOVA Results for Putamen BPnd**

| Effect | Sum Sq | Mean Sq | NumDF | DenDF | F | p (raw) | p (FDR) | partial η² |
|---|---|---|---|---|---|---|---|---|
| Condition | 0.0177 | 0.0088 | 2 | 30.749 | 1.399 | 0.262 | 0.696 | 0.083 |
| Sex | 0.0019 | 0.0019 | 1 | 17.322 | 0.307 | 0.586 | 0.696 | 0.017 |
| Condition × Sex | 0.0070 | 0.0035 | 2 | 30.749 | 0.557 | 0.579 | 0.696 | 0.035 |

**Supplementary Table 29. Pairwise Comparisons for Putamen BPnd**

| Comparison | Estimate | SE | df | t-ratio | Raw p-value | FDR p-value | Hedges' g |
|---|---|---|---|---|---|---|---|
| Female Buffering vs Baseline | -0.085 | 0.088 | 30.0 | -0.964 | 0.343 | 0.570 | -0.176 |
| Female Buffering vs Stress | 0.037 | 0.097 | 31.9 | 0.383 | 0.705 | 0.755 | 0.068 |
| Female Baseline vs Stress | -0.048 | 0.083 | 29.1 | -0.576 | 0.569 | 0.657 | -0.107 |
| Male Buffering vs Baseline | -0.163 | 0.081 | 27.2 | -2.023 | 0.053 | 0.138 | -0.388 |
| Male Buffering vs Stress | 0.209 | 0.076 | 27.6 | 2.748 | **0.010**** | 0.138 | 0.523 |
| Male Baseline vs Stress | 0.047 | 0.079 | 28.3 | 0.587 | 0.562 | 0.657 | 0.110 |
| Baseline Males vs Females | -0.157 | 0.095 | 39.3 | -1.655 | 0.106 | 0.198 | -0.528 |
| Buffering Males vs Females | -0.234 | 0.104 | 41.8 | -2.245 | **0.030*** | 0.138 | -0.694 |
| Stress Males vs Females | -0.062 | 0.098 | 40.4 | -0.633 | 0.530 | 0.657 | -0.199 |
| Pooled: Baseline vs Buffering | -0.124 | 0.060 | 28.7 | -2.075 | **0.047*** | 0.138 | -0.387 |

| | | | | | | | |
|---|---|---|---|---|---|---|---|
| Pooled: Baseline vs Stress | -0.001 | 0.058 | 28.7 | -0.012 | 0.991 | 0.991 | -0.002 |
| Pooled: Buffering vs Stress | 0.123 | 0.062 | 30.2 | 1.995 | 0.055 | 0.138 | 0.363 |

**Insula**

Neither condition contrast reached statistical significance at the model level. Sex was not a significant predictor (β = –0.063, SE = 0.048, t = –1.30, p = 0.209). Interaction terms were not significant for either buffering × sex (p = 0.526) or stress × sex (p = 0.479). Model $R^2$ values were $R^2$ marginal = 0.151 and $R^2$ conditional = 0.407.

Type II ANOVA revealed no significant main effect of condition (F(2, 30.16) = 2.79, p = 0.076, FDR-corrected p = 0.339, partial $\eta^2$ = 0.156) or sex (F(1, 18.90) = 1.66, p = 0.213, FDR-corrected p = 0.355, partial $\eta^2$ = 0.081). The condition × sex interaction was also not significant (F(2, 30.56) = 0.93, p = 0.405, FDR-corrected p = 0.506, partial $\eta^2$ = 0.057). As there were no significant main effects or interactions, pairwise comparisons are not reported.

**Supplementary Table 30. Linear Mixed Model (LMM) of Insula BPnd**

| Predictor | Estimate | Std. Error | df | t value | p-value |
|---|---|---|---|---|---|
| Intercept | 1.913 | 0.048 | 19.020 | 39.703 | < 0.001 |
| Buffering vs Baseline | –0.088 | 0.047 | 29.562 | –1.883 | 0.070 |
| Stress vs Baseline | 0.096 | 0.050 | 31.100 | 1.930 | 0.063 |
| Sex (Male) | –0.063 | 0.048 | 19.020 | –1.300 | 0.209 |
| Buffering × Male | –0.030 | 0.047 | 29.562 | –0.641 | 0.526 |
| Stress × Male | –0.036 | 0.050 | 31.100 | –0.717 | 0.479 |

**Supplementary Table 31. Type II ANOVA Results for Insula BPnd**

| Effect | Sum Sq | Mean Sq | NumDF | DenDF | F | p (raw) | p (FDR) | partial $\eta^2$ |
|---|---|---|---|---|---|---|---|---|
| Condition | 0.2977 | 0.1488 | 2 | 30.157 | 2.7861 | 0.076 | 0.339 | 0.156 |
| Sex | 0.0886 | 0.0886 | 1 | 18.902 | 1.6594 | 0.213 | 0.355 | 0.081 |
| Condition × Sex | 0.0994 | 0.0497 | 2 | 30.567 | 0.9310 | 0.405 | 0.506 | 0.057 |

**Ventral Pallidum**

Neither condition contrast reached significance: buffering did not differ from baseline (β = –0.039, SE = 0.045, t = –0.87, p = 0.391), and stress did not differ from baseline (β = 0.057, SE =

0.047, t = 1.20, p = 0.238). Sex was also not a significant predictor (β = –0.055, SE = 0.042, t = –1.32, p = 0.205). The condition × sex interactions were not significant for buffering (p = 0.568) or stress (p = 0.416). Model $R^2$ values were $R^2$ marginal = 0.087 and $R^2$ conditional = 0.284.

A Type II ANOVA revealed no significant main effect of condition (F(2, 27.21) = 0.68, p = 0.516, FDR-corrected p = 0.515, partial $\eta^2$ = 0.047) or sex (F(1, 15.19) = 1.91, p = 0.187, FDR-corrected p = 0.187, partial $\eta^2$ = 0.11). The condition × sex interaction was also not significant (F(2, 27.75) = 0.36, p = 0.703, FDR-corrected p = 0.703, partial $\eta^2$ = 0.026). As there were no significant main effects or interactions, pairwise comparisons are not reported.

**Supplementary Table 32. Linear Mixed Model (LMM) of Ventral Pallidum BPnd**

| Predictor | Estimate | Std. Error | df | t value | p-value |
|---|---|---|---|---|---|
| Intercept | 1.414 | 0.042 | 15.297 | 33.904 | < 0.001 |
| Buffering vs Baseline | –0.039 | 0.045 | 26.485 | –0.872 | 0.391 |
| Stress vs Baseline | 0.057 | 0.047 | 28.464 | 1.204 | 0.238 |
| Sex (Male) | –0.055 | 0.042 | 15.297 | –1.323 | 0.205 |
| Buffering × Male | –0.026 | 0.045 | 26.485 | –0.579 | 0.568 |
| Stress × Male | 0.039 | 0.047 | 28.464 | 0.825 | 0.416 |

**Supplementary Table 33. Type II ANOVA Results for Ventral Pallidum BPnd**

| Effect | Sum Sq | Mean Sq | NumDF | DenDF | F | p (raw) | p (FDR) | partial $\eta^2$ |
|---|---|---|---|---|---|---|---|---|
| Condition | 0.06686 | 0.03343 | 2 | 27.21 | 0.679 | 0.516 | 0.515 | 0.047 |
| Sex | 0.09423 | 0.09423 | 1 | 15.19 | 1.914 | 0.187 | 0.187 | 0.112 |
| Condition × Sex | 0.03520 | 0.01760 | 2 | 27.75 | 0.358 | 0.703 | 0.703 | 0.026 |

**Parahippocampal Gyrus**

Neither condition contrast reached significance: buffering did not differ from baseline (β = –0.003, SE = 0.025, t = –0.12, p = 0.902), and stress did not differ from baseline (β = 0.021, SE = 0.027, t = 0.80, p = 0.430). Sex was not a significant predictor (β = 0.002, SE = 0.029, t = 0.08, p = 0.937). The buffering × sex interaction was non-significant (p = 0.190), although the stress × sex interaction reached significance (β = –0.070, SE = 0.027, t = –2.62, p = 0.014). Model $R^2$ values were $R^2$ marginal = 0.101 and $R^2$ conditional = 0.472.

A Type III ANOVA indicated no significant main effect of condition (F(2, 30.36) = 0.37, p = 0.693, FDR p = 0.878, partial $\eta^2$ = 0.024) or sex (F(1, 19.67) = 0.006, p = 0.937, FDR p = 0.937, partial $\eta^2 \approx 0.0003$). The condition × sex interaction was significant ($F(2, 30.36) = 3.44$, **$p = 0.045$***, FDR $p = 0.225$, partial $\eta^2 = 0.185$).

Post-hoc comparisons showed that none of the within-sex condition contrasts remained significant after FDR correction. In females, effect sizes were small (g range = –0.16 to 0.22). In males, the baseline–buffering (estimate = –0.128, p = 0.044, FDR p = 0.225, g = –0.41) and buffering–stress contrasts (estimate = 0.146, p = 0.017, FDR p = 0.225, g = 0.49) were significant before correction but not after.

Post-hoc pairwise comparisons showed that, after FDR correction, no contrast remained statistically significant. Significant pairwise contrasts prior to FDR correction are reported in Supplementary Table 36.

**Supplementary Table 34. Linear Mixed Model (LMM) of Parahippocampal Gyrus BPnd**

| Predictor | Estimate | Std. Error | df | t value | p-value |
|---|---|---|---|---|---|
| (Intercept) | 0.918 | 0.029 | 19.668 | 31.139 | <0.001 |
| Buffering vs Baseline | –0.003 | 0.025 | 29.568 | –0.124 | 0.902 |
| Stress vs Baseline | 0.021 | 0.027 | 30.782 | 0.800 | 0.430 |
| Male vs Female | 0.002 | 0.029 | 19.668 | 0.080 | 0.937 |
| Buffering × Male | 0.033 | 0.025 | 29.568 | 1.342 | 0.190 |
| Stress × Male | –0.070 | 0.027 | 30.782 | –2.622 | **0.014** |

**Supplementary Table 35. Type III ANOVA Results for Parahippocampal Gyrus BPnd**

| Effect | Sum Sq | Mean Sq | NumDF | DenDF | F | p (raw) | p (FDR) | partial $\eta^2$ |
|---|---|---|---|---|---|---|---|---|
| Condition | 0.011 | 0.006 | 2 | 30.364 | 0.372 | 0.693 | 0.878 | 0.024 |
| Sex | 0.000 | 0.000 | 1 | 19.668 | 0.006 | 0.937 | 0.937 | ~0.000 |
| Condition × Sex | 0.104 | 0.052 | 2 | 30.364 | 3.439 | **0.045*** | 0.225 | 0.185 |

**Supplementary Table 36: Pairwise Comparisons for Parahippocampal Gyrus BPnd**

| Comparison | Estimate | SE | df | t-ratio | Raw p-value | FDR p-value | Hedges' g |
|---|---|---|---|---|---|---|---|
| Female Buffering vs Baseline | 0.079 | 0.067 | 29.4 | 1.185 | 0.245 | 0.670 | 0.218 |
| Female Buffering vs Stress | –0.067 | 0.074 | 31.0 | –0.910 | 0.370 | 0.670 | –0.163 |

| | | | | | | | |
|---|---|---|---|---|---|---|---|
| Female Baseline vs Stress | 0.012 | 0.063 | 28.7 | 0.192 | 0.849 | 0.910 | 0.036 |
| Male Buffering vs Baseline | –0.128 | 0.061 | 26.9 | –2.116 | **0.044*** | 0.225 | –0.408 |
| Male Buffering vs Stress | 0.146 | 0.057 | 27.4 | 2.553 | **0.017*** | 0.225 | 0.488 |
| Male Baseline vs Stress | 0.018 | 0.060 | 27.9 | 0.307 | 0.761 | 0.878 | 0.058 |
| Baseline Males vs Females | 0.072 | 0.076 | 36.9 | 0.948 | 0.349 | 0.670 | 0.312 |
| Buffering Males vs Females | –0.136 | 0.083 | 40.4 | –1.639 | 0.109 | 0.409 | –0.516 |
| Stress Males vs Females | 0.078 | 0.078 | 38.3 | 0.999 | 0.324 | 0.670 | 0.323 |
| Pooled: Baseline vs Buffering | –0.025 | 0.045 | 28.3 | –0.543 | 0.591 | 0.878 | –0.102 |
| Pooled: Baseline vs Stress | 0.015 | 0.043 | 28.3 | 0.351 | 0.728 | 0.878 | 0.066 |
| Pooled: Buffering vs Stress | 0.040 | 0.047 | 29.6 | 0.850 | 0.402 | 0.670 | 0.156 |
